# Actomyosin activity-dependent apical targeting of Rab11 vesicles reinforces apical constriction

**DOI:** 10.1101/2021.03.07.434309

**Authors:** Wei Chen, Bing He

## Abstract

During tissue morphogenesis, cell shape changes resulting from cell-generated forces often require active regulation of intracellular trafficking. How mechanical stimuli influence intracellular trafficking and how such regulation impacts tissue mechanics are not fully understood. In this study, we identify an actomyosin dependent mechanism involving Rab11- mediated trafficking in regulating apical constriction in the *Drosophila* embryo. During *Drosophila* mesoderm invagination, apical actin and Myosin II (actomyosin) contractility induces apical accumulation of Rab11-marked vesicle-like structures (“Rab11 vesicles”) by promoting a directional bias in dynein mediated vesicle transport. At the apical domain, Rab11 vesicles are enriched near the adherens junctions (AJs). The apical accumulation of Rab11 vesicles is essential to prevent fragmented apical AJs, breaks in the supracellular actomyosin network and a reduction in the apical constriction rate. This Rab11 function is separate from its role in promoting apical Myosin II accumulation. These findings suggest a feedback mechanism between actomyosin activity and Rab11-mediated intracellular trafficking that regulates the force generation machinery during tissue folding.

## Introduction

Cell-generated mechanical forces play a central role in tissue morphogenesis. Genetically prescribed cellular forces can drive cell shape change and cell motion as a direct physical outcome, thereby mediating spatially and temporally defined tissue deformation (Collinet and Lecuit, 2021; Gilmour et al., 2017). In addition, mechanical forces can impact morphogenesis by triggering various cellular activities ranging from cytoskeleton reorganization to changes in gene expression (Fletcher and Mullins, 2010; Kirby and Lammerding, 2016; Sun and Irvine, 2016; Uhler and Shivashankar, 2017). Such active processes allow cells and tissues to respond to mechanical inputs adaptively and thereby increase the robustness of tissue morphogenesis. However, our knowledge of the active response of cells to mechanical forces remains limited.

In this work, we used mesoderm invagination during *Drosophila* gastrulation as a model to study how cells respond to contractile forces generated by actomyosin networks. Immediately before gastrulation, *Drosophila* embryos undergo an atypical cleavage called cellularization, during which peripherally localized syncytial nuclei are partitioned into a monolayer of epithelial cells (Mazumdar and Mazumdar, 2002). Mesoderm invagination is initiated by apical constriction resulted from graded apical activation of actomyosin in a 12-cell wide region in the ventrally localized mesodermal primordium (Heer et al., 2017; Leptin and Grunewald, 1990; Lim et al., 2017; Sweeton et al., 1991). These cells subsequently invaginate as the epithelium bends inwards, resulting in the formation of a furrow on the ventral side of the embryo. The upstream signaling pathway that activates apical constriction has been well characterized (reviewed in Gheisari et al., 2020; Gilmour et al., 2017; Martin, 2020). In response to the dorsal-ventral patterning information, the prospective mesodermal cells express two transcription factors, Twist and Snail, which in turn trigger the apical recruitment and activation of RhoGEF2 through GPCR signaling (Jha et al., 2018; Kerridge et al., 2016; Manning et al., 2013). RhoGEF2 further activates the small GTPase Rho1 (the *Drosophila* homologue of RhoA) and leads to the activation of Myosin II through Rho1’s effector Rho-associated protein kinase (Rok). Activated Myosin II forms an supracellular actomyosin network at the apical cortex that is interconnected across the tissue through apical adherens junctions (AJs) (Amano et al., 1996; Dawes-Hoang et al., 2005; Kimura et al., 1996; Mason et al., 2013; Nikolaidou and Barrett, 2004; Vasquez et al., 2014; Winter et al., 2001). Within individual cells, the contractions of apical actomyosin network pull AJ sites inwards, causing progressive apical cell area reduction (Martin and Goldstein, 2014; Martin et al., 2009; Martin et al., 2010; Mason et al., 2013).

Apical constriction can drive reorganization from subcellular to tissue levels through both active and passive means. Apical constriction drives a tissue-scale viscous flow at the tissue interior (He et al., 2014). Certain organelles, such as the nucleus, appear to move passively with the flow by advection (Gelbart et al., 2012). Other subcellular structures, in contrast, undergo active reorganization in response to apical forces. For examples, apical actomyosin contractility has been shown to mediate apical shift and stabilization of the AJs and promote the formation of a medioapically-localized noncentrosomal microtubule-organization center (Weng and Wieschaus, 2016; Ko et al., 2019). It remains unclear whether other subcellular structures also undergo active remodeling in response to apical constriction.

Rab11, a small GTPase of the Rab family, is a well-established marker for recycling endosomes (Welz et al., 2014). Like typical small GTPases, Rab11 switches between active and inactive states depending on the phosphorylation state of its bound guanine nucleotide. In addition to recycling endosomes, Rab11 has also been reported to localize to the trans-Golgi network (TGN) and post-Golgi vesicles. It functions in both exocytic and endocytic recycling pathways by regulating vesicular transport from TGN and recycling endosomes to the plasma membrane (Benli et al., 1996; Chen et al., 1998; Jedd et al., 1997; Pelissier et al., 2003; Takahashi et al., 2012; Ullrich et al., 1996; Welz et al., 2014). Rab11 regulates multiple vesicular trafficking steps, including vesicle formation, transport and tethering through its various downstream effectors. For example, Rab11 directly interacts with the actin motor Myosin V (Lapierre et al., 2001; Lipatova et al., 2008). It can also form complexes with microtubule motors dynein and kinesins through adaptor proteins (Welz et al., 2014). These interactions allow Rab11 vesicles to be transported along both actin and microtubule filaments (Horgan et al., 2010; Schonteich et al., 2008; Schuh, 2011; Wang et al., 2008).

Here, we present evidence that apical actomyosin contractility induces biased transport of Rab11-positive vesicles towards the cell apex in a dynein- and microtubule-dependent manner. At the apical side, Rab11 vesicles are enriched at the vicinity of AJs. Eliminating these vesicles through acute inactivation of Rab11 results in defects in AJ organization and breaks in the supracellular actomyosin network. Our findings reveal an intimate interplay between apical actomyosin contractions and Rab11-mediated vesicle trafficking that serves as a feedback mechanism to reinforce the structural integrity of the apical actomyosin network.

## Results

### Rab11 accumulates apically in the constricting cells during ventral furrow formation

In a search for potential changes in subcellular organization induced by apical actomyosin contractions, we found that Rab11-labeled compartments undergo prominent reorganization during apical constriction (Figure 1A-E; Figure S1). Live imaging of embryos expressing endogenously YFP-tagged Rab11 and image segmentation (Berg et al., 2019; Methods) revealed two main types of Rab11-positive structures (Figure 1F, G; Figure S1B): (1) One or a few large perinuclear compartments residing at the apical side of each nucleus (hereafter “perinuclear Rab11 compartment”). These compartments likely represented the trafficking intermediate described in a previous study based on similarities in morphology and localization (Pelissier et al., 2003). And (2) Small punctate structures. These puncta were less than 1 μm in diameter and appeared as diffraction limited spots. Based on previous literature, these Rab11 puncta could be small transport vesicles, recycling endosomes and/or other type of transport carrier (Welz et al., 2014) (“Rab11 vesicles” herein). Before gastrulation, Rab11 vesicles were present near the basal side of the perinuclear Rab11 compartments (Figure 1C, 0 min; Figure 1F, G, 0 min). As the cells constricted apically, perinuclear Rab11 compartments moved basally following the basal movement of the nuclei (Figure 1C, blue outlines, 9 min; Figure 1F, G, 7 min). Meanwhile, Rab11 vesicles appeared near the cell apices and accumulated apically (Figure 1C-G). The difference in Rab11 vesicle distribution before and after apical constriction was further confirmed by quantification of Rab11 vesicle intensities at various apicobasal positions (Figure 1H; Methods). The Rab11 vesicles were highly dynamic, yet at any given time the majority accumulated either adjacent to or slightly basal to the apical actomyosin network (Figure 1E). No apical accumulation of Rab11 vesicles was observed in the surrounding ectodermal cells that did not undergo apical constriction (Figure 1C; Figure S1A). In addition, we did not observe obvious apical enrichment in the constricting cells for the other organelles we examined (Figure S1C-H).

**Figure 1.**
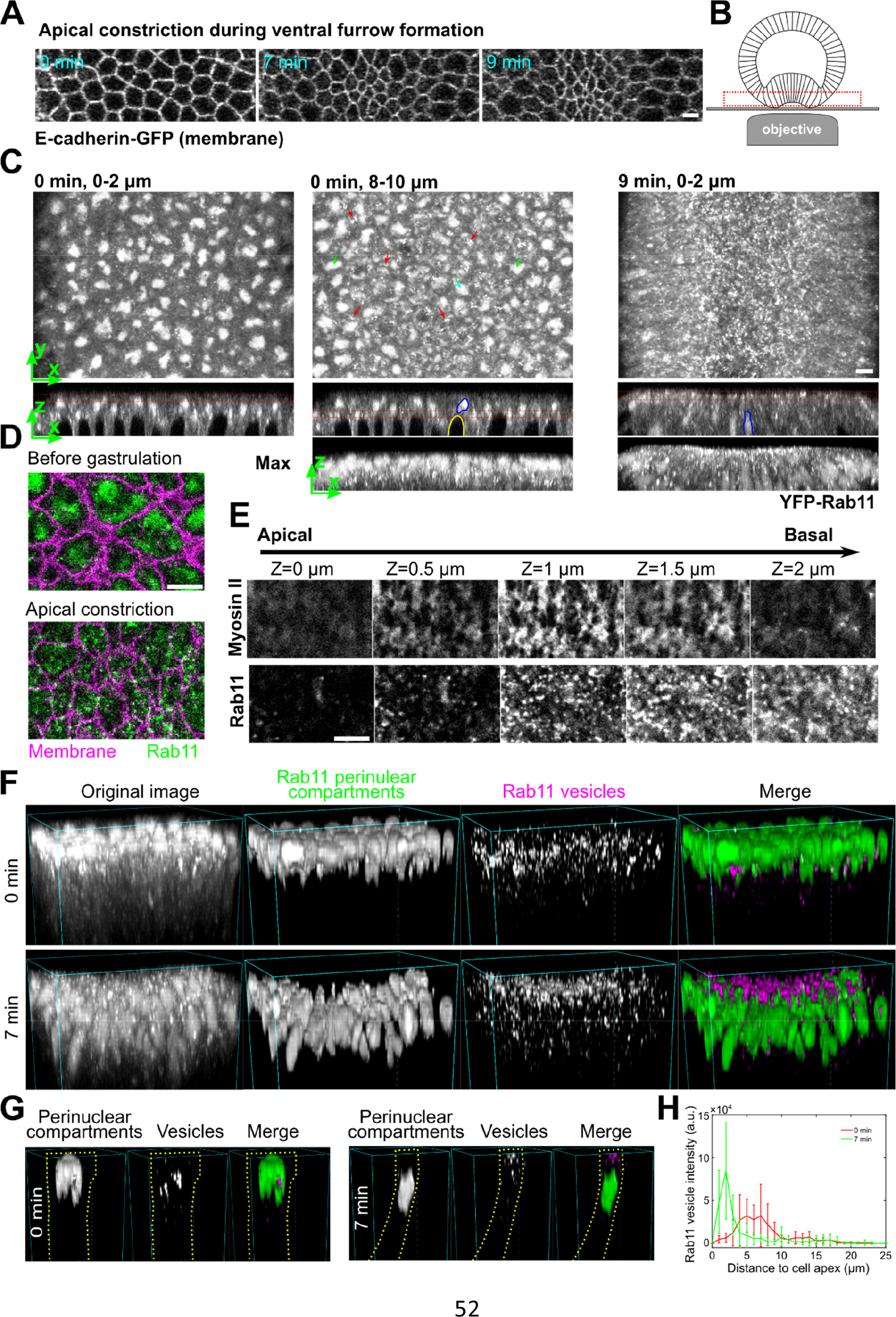
Rab11-positive vesicle-like structures accumulate apically in constricting cells during ventral furrow formation. **(A)** Apical constriction during ventral furrow formation. Scale bar, 10 μm. **(B)** The imaging setup for acquiring the en face view of constricting cells. **(C)** Rab11 compartments (endogenous YFP-Rab11) in an embryo at the onset (0 min) and 9 min into ventral furrow formation. Top panel, maximum projection over 2 μm depth from en face view; middle and bottom panels, a single slice and maximum projection of the cross-section view, respectively. Red boxes: Z position of the corresponding en face view. Red arrows: perinuclear Rab11 vesicles; green arrows and blue outlines: perinuclear Rab11 compartments; yellow outline: nucleus. **(D)** Zoom-in view of a small ROI in the ventral region of the embryo. **(E)** Montage showing Rab11 and Myosin II distribution over depth during apical constriction. **(F, G)** 3D reconstruction of Rab11 positive structures in ventral mesoderm constricting cells. 0 min: onset of gastrulation. Rab11 perinuclear compartments and vesicles are segmented separately using ilastik. An individual cell is shown in G as an example. Cell outlines (yellow dotted lines) are determined based on Gap43-mCherry signal. **(H)** The intensity profile of summed intensity of Rab11 vesicles along Z direction. 0 μm: cell apex. Error bar: s.d. N = 13 cells. All scale bars are 5 μm unless mentioned otherwise.

### The apical enrichment of Rab11 vesicles could not be readily explained by cytoplasmic advection and suggests the involvement of active regulation

Apical accumulation of Rab11 vesicles depends on apical actomyosin network but does not require intact adherens junctions or apical shrinking The spatiotemporal correlation between the apical Myosin II activation and the apical accumulation of Rab11 vesicles prompted us to examine the relationship between the two processes. To test whether apical activation of Myosin II is required for apical accumulation of Rab11 vesicles, we injected embryos with the Rok inhibitor Y-27632 (Narumiya et al., 2000). Injection of Y-27632 at the end of cellularization completely prevented apical Myosin II activation and abolished apical Rab11 vesicle accumulation (Figure 2A). Next, we adapted a previously described on-stage injection protocol (Coravos and Martin, 2016) to test the effect of Y-27632 injection on Rab11 vesicles after they accumulated apically. After drug injection, Myosin II immediately (< 1 min) dissociated from the apical cortex (Figure 2B, B’). About 4 minutes after injection, Rab11 vesicles were largely depleted from the apical surface (Figure 2B). The morphology of perinuclear Rab11 compartments was not obviously affected by Y-27632 injection (Figure 2B, 3 – 6 μm). Consistent with the Y-27632 treatment, knockdown of the myosin heavy chain Zipper also inhibited apical Rab11 vesicle accumulation (Figure 2C). Together, these observations indicate that the induction and maintenance of apical accumulation of Rab11 vesicles depends on the recruitment and/or activation of apical Myosin II. In further support of this notion, we found that an expansion of the actomyosin activation domain in ventralized *Spn27A* mutant embryos resulted in a similar expansion of the region with apical accumulation of Rab11 vesicles (Figure 2D; Ligoxygakis et al., 2003).

**Figure 2.**
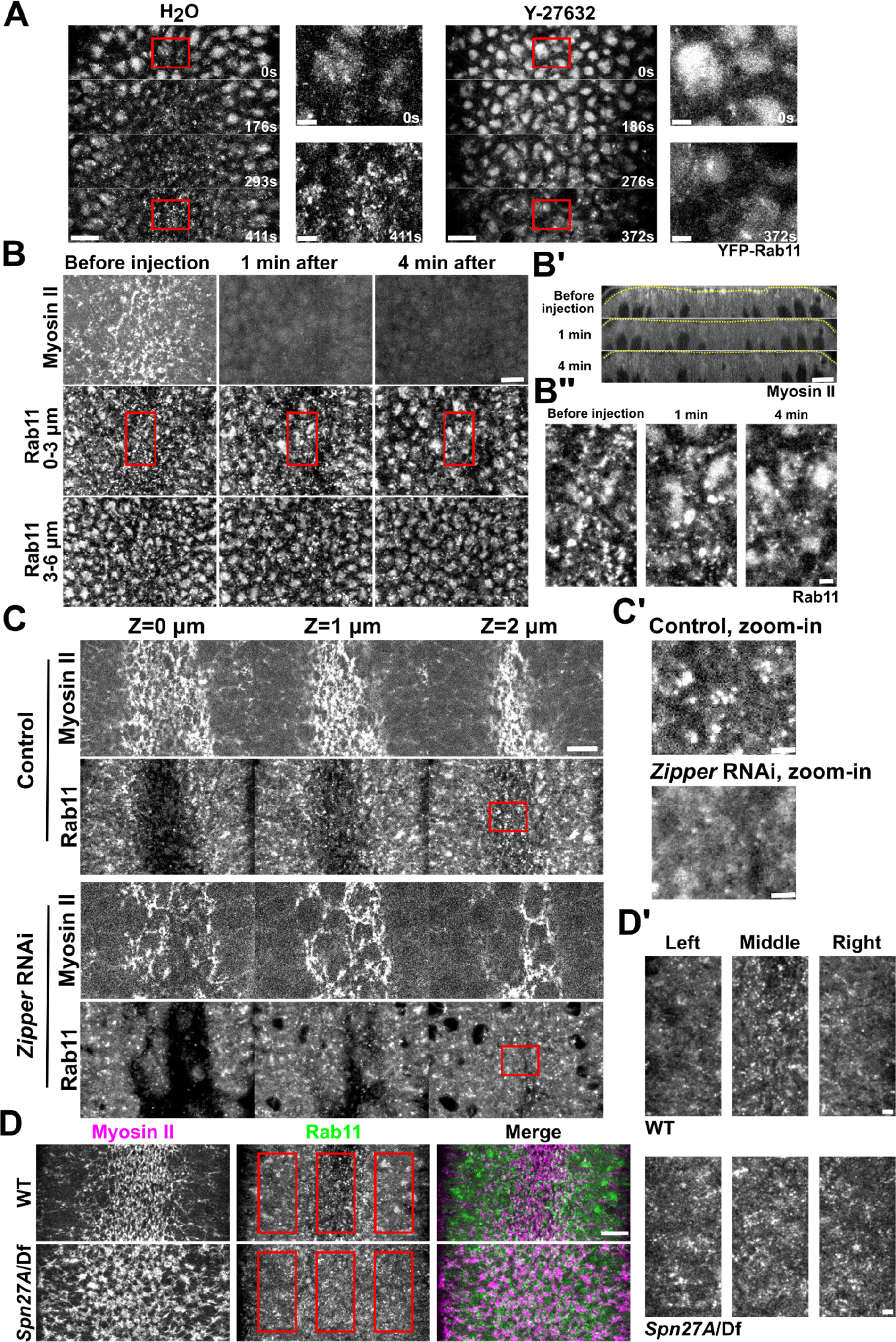
Apical accumulation of Rab11 vesicles depends on apical Myosin II activation. **(A)** Inhibition of Myosin II activation by Y-27632 injection at late cellularization prevents apical Rab11 vesicle accumulation. (A’) Zoom-in view of the outlined region in A. 0 sec: onset of gastrulation. Y-27632 injection: N = 4 embryos; H2O injection: N = 3 embryos. **(B)** Injection of Y-27632 during apical constriction causes rapid diminishing of apical Rab11 vesicles. N = 4 embryos. (B’) Cross-section view. Yellow lines: apical surface. (B’’) Zoom-in view of B. **(C)** Knockdown of myosin heavy chain Zipper inhibits apical accumulation of Rab11 vesicle. N = 4 embryos for each genotype. (C’) Zoom-in view of C. **(D)** *Spn27A*/Df mutant embryos exhibit ventralized phenotype with expanded domain of apical Myosin II activation and apical Rab11 vesicle accumulation (N = 3 embryos). (D’) Zoom-in view of D. Images were Gaussian filtered with a radius of 0.5 pixel. Fewer z-slices were projected for the zoom-in view to improve the signal/noise. All scale bars for zoom-in view, 2 μm; scale bars for the rest, 10 μm. All en face views are maximum projections of a ∼3 – 4 μm-stack below apical surface of the constriction domain.

**Figure 3.**
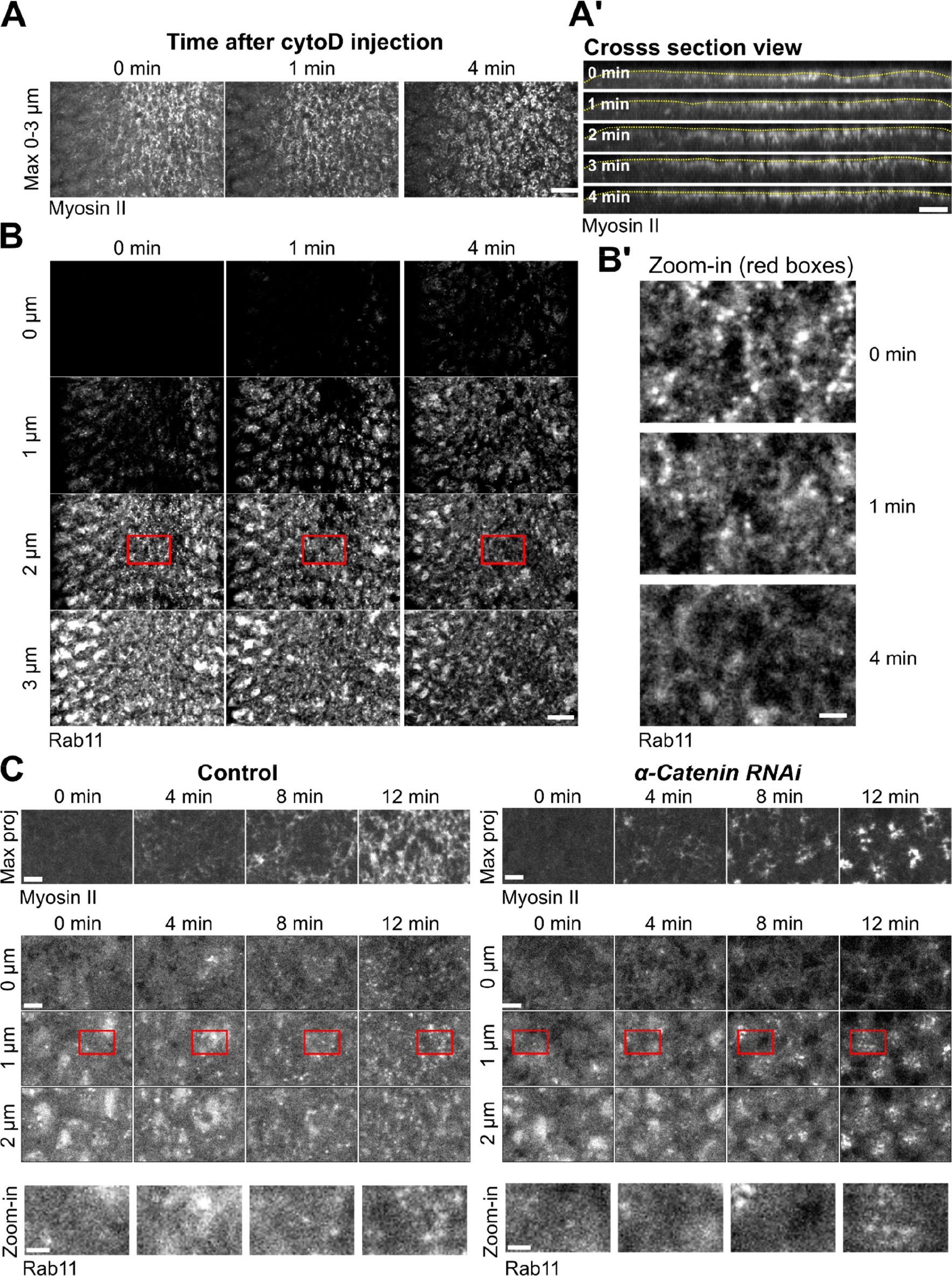
Apical accumulation of Rab11 vesicles depends on F-actin but not does not require intact AJs. **(A, A’, B, B’)** Cytochalasin D injection (0 min) during apical constriction results in rapid diminishing of apical Rab11 vesicles without causing immediate apical myosin loss. (A) Maximum projection of apical Myosin II. Scale bar, 10 μm. (A’) Cross-section view. Yellow lines: apical surface. Scale bar, 10 μm. (B) Enface view showing apical Rab11 vesicles. Scale bar, 10 μm. (B’) Zoom-in view of B. Scale bar, 2 μm. **(C)** Knockdown of junctional component α-Catenin does not prevent apical accumulation of Rab11 vesicles. Scale bars, 5 μm. Scale bars for zoom-in view, 2 μm.

Since cortical actin network is required for Myosin II to generate contractile forces (Martin et al., 2009), we asked whether apical Rab11 vesicle accumulation depends on F-actin. To this end, we injected embryos with cytochalasin D (cytoD) to acutely disrupt actin network during apical constriction. Upon cytoD injection, the apical side of the constricted cell immediately relaxed, suggesting a rapid loss of myosin contractility (Figure 3A; Video 1).

Interestingly, Myosin II remained enriched apically in the first four minutes after injection (Figure 3A). In contrast, apical Rab11 vesicles quickly diminished after injection, similar to what we observed in Y-27632 injected embryos (Figure 3B). This result indicates that the accumulation of Myosin II at the apical domain is not sufficient to induce apical Rab11 vesicle accumulation without F-actin. Together, our observations so far suggest that an intact, contractile apical actomyosin network is critical for apical accumulation of Rab11 vesicles.

During ventral furrow formation, contractile forces generated by apical actomyosin network result in progressive apical shrinking in the presumptive mesoderm and a buildup of apical tension across the constriction domain, both require apical AJs (Martin et al., 2009; Martin et al., 2010; Mason et al., 2013). We therefore asked whether apical shrinking or the presence of intact AJs is required for apical accumulation of Rab11 vesicles. We found that knockdown of α-Catenin, a core component of AJ complex, did not prevent Rab11 vesicle accumulation, although this treatment impaired the connection between the actomyosin network and the cell membrane, causing Myosin II to coalesce into clustered spots at the center of the medioapical domain (Martin et al., 2010; Figure 3C). In addition, we found that induction of apical actomyosin contractility in dorsal cells without accompanying apical shrinking through Fog overexpression (Morize et al., 1998; Methods) is sufficient to induce apical accumulation of Rab11 vesicles (Figure 4A-C). Finally, we found that apical shrinking during dorsal fold formation, a Myosin II-independent tissue folding process (Wang et al., 2012), did not induce apical accumulation of Rab11 vesicles (Figure 4D, E). Taken together, these results indicate that the apical accumulation of Rab11 vesicles, which we have shown requires apical contractile actomyosin network, does not depend on mesodermal cell fate, apical shrinking or integration of tensile forces across the tissue.

**Figure 4.**
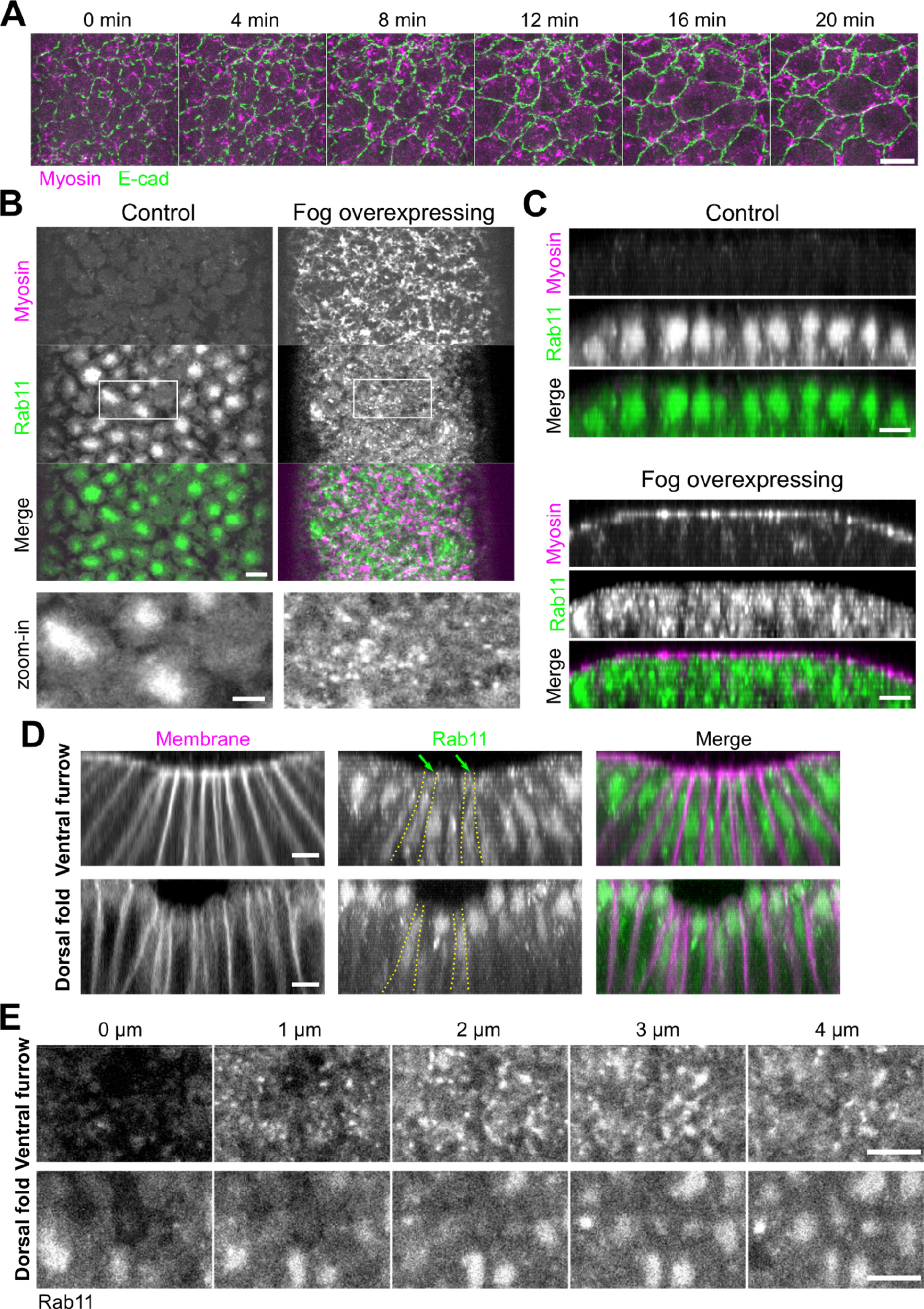
Apical actomyosin activation, but not apical cell area reduction, is critical for apical accumulation of Rab11 vesicles. **(A)** Ectopic expression of Fog results in Myosin II activation in dorsal cells but does not cause apical area reduction. N = 3 embryos. Scale bar, 10 μm. **(B, C)** Ectopic activation of Myosin II in dorsal cells by overexpression of Fog induces ectopic apical accumulation of Rab11 vesicles. (B) Enface view. (C) Cross section view. Scale bars, 5 μm. Scale bar for zoom-in view, 2 μm. **(D, E)** Comparison of YFP-Rab11 in ventral furrow and dorsal fold. Yellow outlines: example cells displaying apical area reduction. Dorsal fold cells do not show apical Rab11 vesicle accumulation despite undergoing apical shrinking. (D) and (E): Cross-section view and en face view, respectively. Scale bar, 5 μm. Green arrows: Rab11 vesicles. Scale bar, 5 μm.

### Rab11 vesicles undergo biased transport towards the cell apex during apical constriction

We next examined how Rab11 vesicles accumulate at the apical surface during apical constriction. Apical endocytosis has been shown to play an important role in apical constriction as the apical membrane shrinks over time (Lee and Harland, 2010; Mateus et al., 2011; Miao et al., 2019). Rab11 vesicles may be derived from endocytosis of apical membranes. However, when we chased endocytosed cell membrane using FM4-64, a lipophilic dye (Methods), we barely detected any colocalization between FM4-64 positive vesicles and Rab11 vesicles, although a majority of FM4-64 positive vesicles colocalized with Rab5-marked early endosomes (Figure S2A-C, Video 2). Consistent with this observation, we found that blocking endocytosis using the temperature sensitive mutant of *shibire* (*shi^ts^*), the *Drosophila* homolog of dynamin (van der Bliek and Meyerowrtz, 1991), did not prevent apical accumulation of Rab11 vesicles (Figure S2D, E). Together, these results indicate that endocytosed apical membrane is not the major source of apical Rab11 vesicles.

An intriguing possibility is that the apical Rab11 vesicles are supplied from the more basal side of the cell by directional transport. To capture the rapid movement of Rab11 vesicles in the apical-basal direction, we focused on the cells located near the medial-lateral boundaries of the constricting domain. These cells are tilted towards the ventral midline at the late stage of apical constriction, making it possible to follow the apical-basal movement of Rab11 vesicles by fast imaging of a single focal plane (Figure 5A). We always selected focal planes where myosin could be observed at the medial side of the cells to ensure that the cells we analyzed were accumulating myosin apically (Figure 5B). Using this approach, we observed constant transport of Rab11 vesicles in the axial direction of the cells (Figure 5B). There were both apically and basally directed movements, with a bias towards the apical direction.

**Figure 5.**
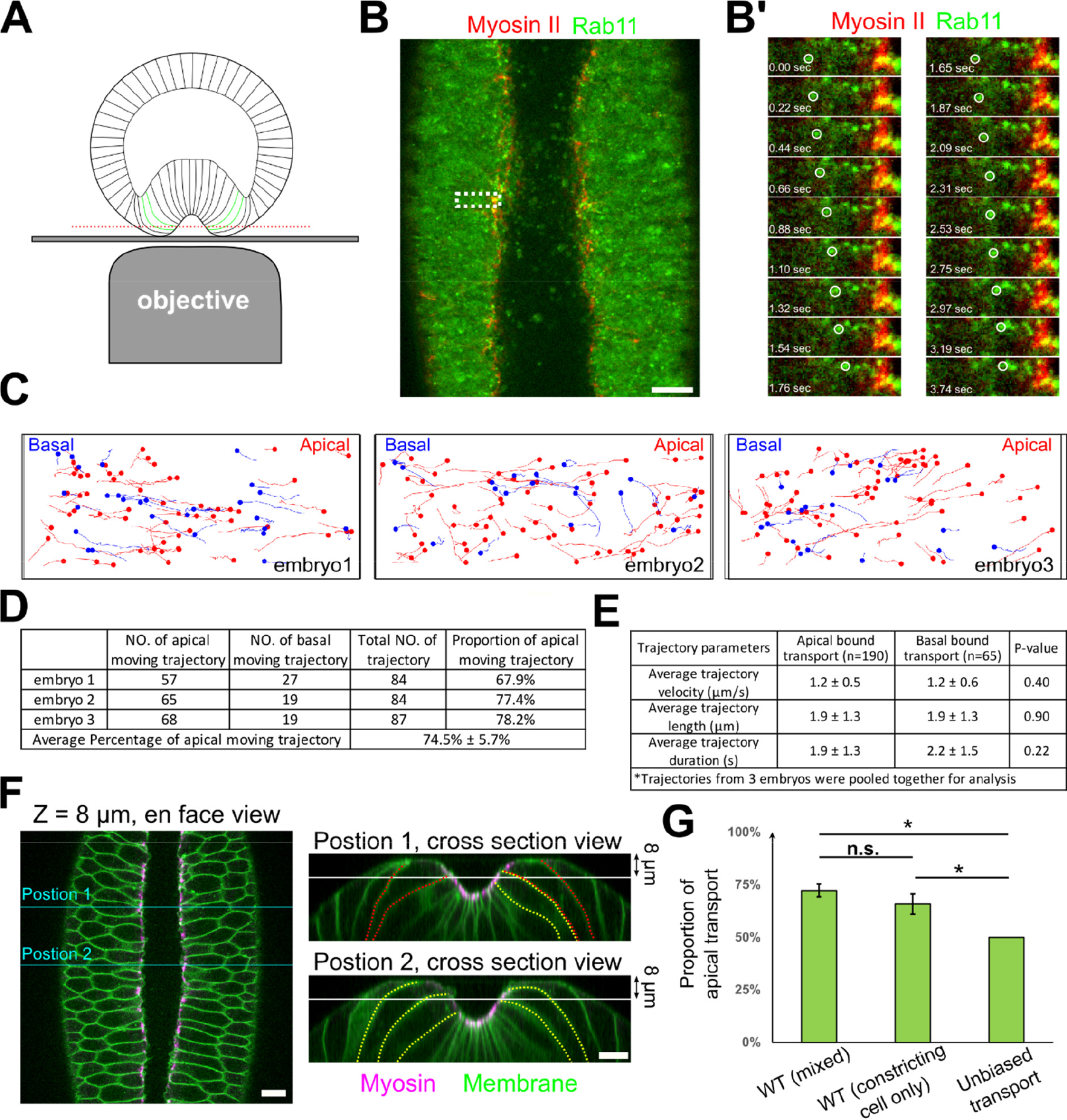
Rab11 vesicles are transported apical-basally with a strong bias in the apical direction. **(A)** Imaging configuration to capture the movement of Rab11 vesicles along the apical-basal direction. Red line: imaging plane. Green outlines: constricting cells being imaged. **(B, B’)** Surface view of an embryo imaged as depicted in A. Scale bar, 10 μm. (B’): zoom-in view of B showing the apical movement of two Rab11 vesicles (white circles). Scale bar, 2 μm. **(C)** Vesicle trajectories in 3 wildtype embryos in a ∼35-sec time window. Red and blue mark apical- and basal-bound trajectories, respectively. **(D)** Counts of apically and basally oriented trajectories. **(E)** Comparison of average trajectory length, velocity and duration (mean ± s.d.) shows no significant difference between apically and basally directed transport. Two tailed Student’s t-test; N = 3 embryos. **(F)** Left: Enface view of an embryo expressing both myosin and membrane markers. Right: Cross-section view of the same embryo at the indicated A-P positions (cyan lines). White line: imaging plane; yellow and red outlines: cells with or without obvious apical shrinking, respectively. Scale bars, 10 μm. **(G)** Rab11 vesicles exhibit directional bias towards the apical side. WT (cells with apical Myosin II activation): quantification of movies with Myosin II and Rab11 markers. The ROI may include a mixture of apically constricted and non-constricted cells that exhibited apical myosin accumulation. WT (constricting cell only): quantification of movies with membrane and Rab11 markers. Only apically constricted cells are quantified (Methods). Two tailed one- sample t-test against 50% was used for comparison to hypothetical unbiased transport. Two tailed Student’s t test was used for comparison between two WT samples. N = 3 embryos for each category. For all panels, error bar: s.d.; ***: p < 0.001; **: p < 0.01; *: p < 0.05; n.s.: p >0.05.

Vesicle tracking revealed that the apically directed movement accounted for approximately three quarters of the total transport events (74.5% ± 5.7%; n = 255 trajectories from 3 embryos, Figure 5C, D, G). Other than the bias in directionality, there was no significant difference in other aspects of the trajectories between apical and basal transport, including the average velocity, the duration and the travel distance of the tractable trajectories (Figure 5E). To further validate the results, we imaged embryos expressing a membrane marker and mCherry-Rab11 (overexpression), which allowed us to quantify transport events only in cells undergoing apical constriction (Figure 5F, Methods). There was no significant difference between the results from the two analyses (Figure 5G). Note that due to our imaging configuration, our analysis of vesicle transport was limited to the apical portion of the cells (typically above the nuclei) that bent over towards the midline. It is unclear whether Rab11 vesicle transport also exists at the more basal region of the cell. Taken together, our results demonstrate that Rab11 vesicles undergo biased transport to the cell apex in cells with active apical actomyosin contractions. This biased transport may account for the apical enrichment of Rab11 vesicles in these cells.

### The apical-basal transport of Rab11 vesicles depends on microtubules and dynein

Because the saltatory and directional movement of Rab11 vesicles was reminiscent of microtubule-dependent transport, we asked whether the transport of Rab11 vesicles is mediated by microtubules. By two-color imaging with mCherry-Rab11 and the microtubule marker Jupiter-GFP (Karpova et al., 2006), we found that most Rab11 vesicles were associated with microtubules or microtubule bundles as they moved apically or basally (Figure 6A; Video 3). Furthermore, on-stage injection of colchicine, a microtubule depolymerization drug, in gastrulating embryos resulted in a rapid (less than a minute) loss of microtubules and a sharp reduction in the directional transport events. The remaining transport events only occurred in regions with residual microtubules (Figure 6B; Video 3).

**Figure 6.**
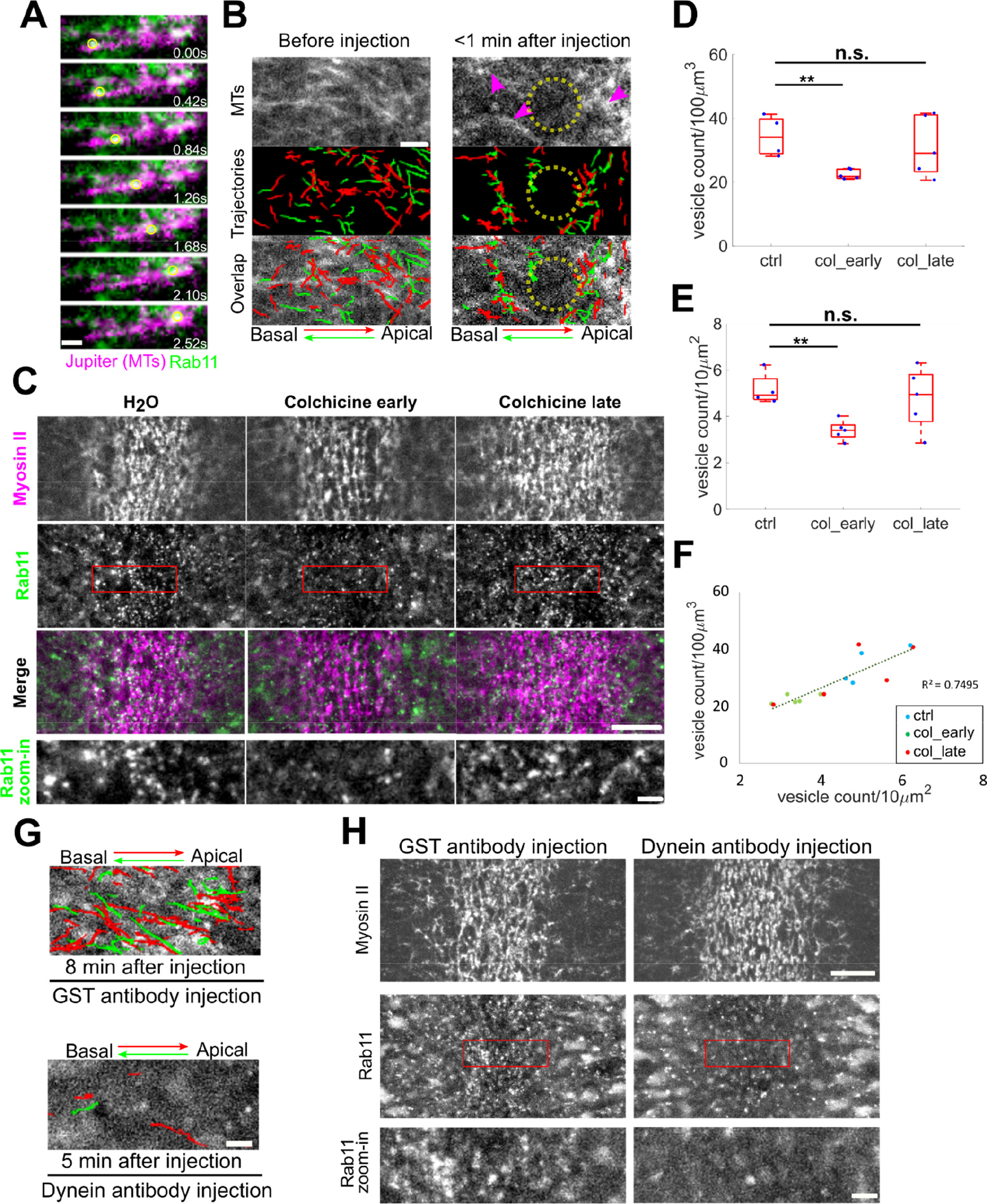
Rab11 vesicle transport requires microtubules and dynein. **(A)** An example of Rab11 vesicle movement (yellow circle) along the microtubules (Jupiter- GFP). Scale bar, 1 μm. **(B)** Acute disruption of microtubules via colchicine injection results in immediate inhibition of Rab11 vesicle transport. Magenta arrowheads: residual microtubules. Yellow circles: region where microtubules were completely disrupted. Scale bar, 2 μm. **(C)** En face view of the constricting domain shortly (< 2 min) after on-stage injection of water or colchicine. Injection of colchicine shortly before but not after the onset of gastrulation (N = 5 embryos for each condition) results in reduced Rab11 vesicle accumulation compared to water injected controls (N = 4 embryos). Scale bar, 10 μm. Bottom panels: zoom-in views. Scale bar, 2 μm. **(D-F)** Quantification of Rab11 vesicle density after water or colchicine injection within a 3.5 μm-stack from apical surface (D) or in a single z plane near apical surface (E) generates consistent results (F). **(G)** Injection of dynein antibody rapidly abolishes both apical (red) and basal (green) transport of Rab11 vesicles. Trajectories show vesicle movement within a 20-sec window at the indicated time after injection. N = 5 embryos for each condition. Scale bar, 2 μm. **(H)** En face view showing inhibition of apical accumulation of Rab11 vesicles upon on-stage injection of dynein antibody immediately before the onset of gastrulation. N = 5 and 6 embryos for GST antibody and dynein antibody injection, respectively. Scale bar, 10 μm. Bottom panels: zoom-in views. Scale bar, 2 μm.

### Together, these results demonstrate that the apical-basal transport of Rab11 vesicles is microtubule dependent

Next, we examined the impact of colchicine injection on the apical accumulation of Rab11 vesicles. In line with the observed role of microtubules in Rab11 vesicle transport, injection of embryos at late cellularization (N = 5 embryos) resulted in a significant reduction in the apical accumulation of Rab11 vesicles compared to control embryos (N = 4 embryos) (Figure 6C-F). The activation of apical Myosin II was normal in the injected embryos, consistent with a recent report (Ko et al., 2019). Unexpectedly, injection of colchicine at or after the onset of gastrulation (N = 5 embryos) resulted in a wider range of phenotypes, and on average there was no significant reduction in the number of apically enriched Rab11 vesicles (Figure 6C-F). We speculate that disruption of microtubules not only reduces the apical-basal transport of Rab11 vesicles but also inhibits the turnover of the vesicles that are already accumulated apically. This combined effect may obscure the effect of loss of microtubules after the onset of gastrulation.

Finally, we tested the role of microtubule motor dynein on the transport of Rab11 vesicles. We found that injection of antibody against the dynein intermediate chain (DIC) (Yi et al., 2011) during apical constriction resulted in a rapid reduction of vesicle transport events in both apical and basal directions, whereas injection of a control GST antibody had no effect (Figure 6G, Video 4). Furthermore, the apical accumulation of Rab11 vesicles was greatly reduced in dynein antibody-injected embryos but not in the control antibody-injected embryos (Figure 6H). We were not able to perform similar test on the role of kinesin motors due to a lack of functional reagents. However, the pronounced impact of dynein inhibition on the transport and apical accumulation of Rab11 vesicles suggests that dynein plays a predominant role in the apical-basal transport of Rab11 vesicles during apical constriction.

The observation that inhibiting dynein abolishes both apical and basal transport of Rab11 vesicles may reflect the bidirectional nature of the microtubule populations in the space between cell apices and nuclei during apical constriction (Ko et al., 2019).

### The apical bias of Rab11 vesicle transport is rapidly abolished upon acute inhibition of apical actomyosin contractility

Because the apical enrichment of Rab11 vesicles depends on actomyosin activity, we asked whether the biased basal-to-apical transport of Rab11 vesicles also depends on actomyosin activity. To address this question, we used a recently developed optogenetic tool to acutely inhibit actomyosin activity through blue light-induced recruitment of a dominant negative form of Rho1 (Rho1DN) to the plasma membrane (“Opto-Rho1DN”, Figure 7A; Guo et al., 2021). Using the same imaging configuration as shown in Figure 5A, we examined the immediate impact of actomyosin inhibition on the transport of Rab11 vesicles. As expected, blue light stimulation resulted in rapid loss of apical Myosin II (within ∼30 seconds; Figure 7B). The loss of apical Myosin II at this stage of ventral furrow formation did not cause furrow relaxation or immediate changes in cell orientation. Strikingly, blue light stimulation resulted in a rapid change in the transport of Rab11 vesicles. Within the first 23 seconds after stimulation, the proportion of apical transport was 64.6% (64.6% ± 4.1%; n = 575 trajectories from 3 embryos), lower than that in the wildtype controls (Figure 7C). This proportion dropped to close to 50% (52.3% ± 2.0%; n = 645 trajectories from 3 embryos) in the next 23- second time window, indicating that the directional bias was completely abolished (Figure 7C). Notably, the loss of directional bias was caused by an increase in the basal transport events instead of a decrease in the apical transport events (Figure 7D, E). Other aspects of the transport are comparable between the two time windows after stimulation (Figure 7F).

**Figure 7.**
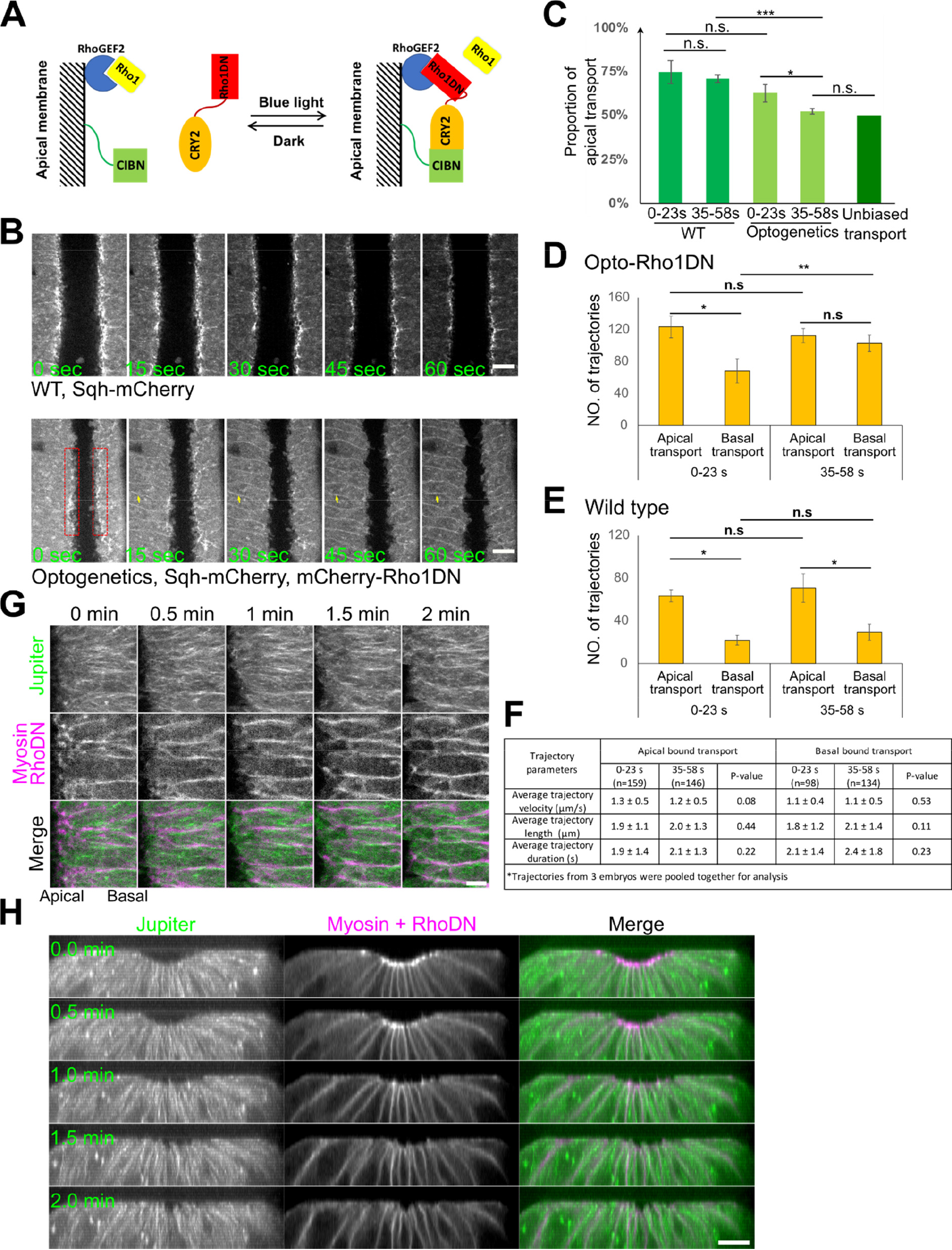
The apical bias of Rab11 vesicle transport depends on activation of apical actomyosin contractility. **(A)** Diagram depicting optogenetic inhibition of Myosin II activation by Opto-Rho1DN. **(B)** Top: upon continuous blue light stimulation, Rho1DN is immediately recruited to the plasma membrane (yellow arrows). Apical myosin II (red boxes) disappears ∼ 30 sec after stimulation. Bottom: A control embryo at similar stage of furrow formation. Scale bar, 10 μm. **(C – F)** The apical bias of the transport is abolished in less than one minute after Opto- Rho1DN stimulation (C, n = 3 embryos). This change is caused by a significant increase in the number of basally directed transport events (D), which is not observed in wildtype embryos (E, n = 3 embryos). There is no significant difference on average trajectory velocity, length and duration of apical and basal bound transports between 0 – 23 sec and 35 – 58 sec post stimulation windows (F, mean ± s.d.). Two tailed one-sample t-test against 50% was used for comparison to hypothetical unbiased transport. Two tailed Student’s t test was used for comparison between conditions. **(G, H)** Microtubules do not exhibit obvious changes in intensity or overall organization in the first two minutes after Opto-Rho1DN stimulation (0 min). (G) En face maximum intensity projection of the apical surface (5 μm thick) showing microtubules (GFP-Jupiter) in an embryo stimulated at the stage comparable to that in B. Scale bar, 5 μm. (H) Cross section view showing an embryo stimulated at a stage slightly earlier than that in (G). Scale bar, 10 μm.

### Therefore, the main effect of actomyosin inhibition on Rab11 vesicle transport is to increase the basal transport events, which results in loss of apical bias of the transport

The observation that the loss of transport bias upon actomyosin inactivation is caused by an increase in the basal transport rather than a reduction in the apical transport suggests that the effect is not due to a general disruption of the microtubule organization. In line with this prediction, we did not observe obvious changes in the density or orientation of microtubules within the first two-minute window after Opto-Rho1DN stimulation (Figure 7G, H). This result is consistent with the previous report that medioapical MTOC only became obviously affected a few minutes after cytoD injection (Ko et al., 2019). Together, these results suggest that apical actomyosin network could impact the directionality of Rab11 vesicle transport partially independent of its role in organizing microtubules in the constricting cells (Ko et al., 2019).

### Rab11 reinforces apical AJs during apical constriction

We next sought to determine the function of apically accumulated Rab11 vesicles. Single plane fast imaging analysis revealed that Rab11 vesicles remained very dynamic after they arrived at the cell apices, with a substantial fraction moving towards apical AJs (Figure 8A; Video 5). The enrichment of Rab11 vesicles near AJs was particularly obvious during the early stage of apical constriction when apical AJs had not yet formed a continuous belt and appeared as discrete foci (Figure 8A). Intensity analysis of E-cadherin-GFP and Rab11- mCherry along cell boundaries revealed a positive correlation between the two intensity profiles (Figure 8B-D; Methods). As a control, similar analysis of Rab11 and a general membrane marker P4M, a PI(4)P binding protein, showed no significant correlation (Figure 8B-D).

**Figure 8.**
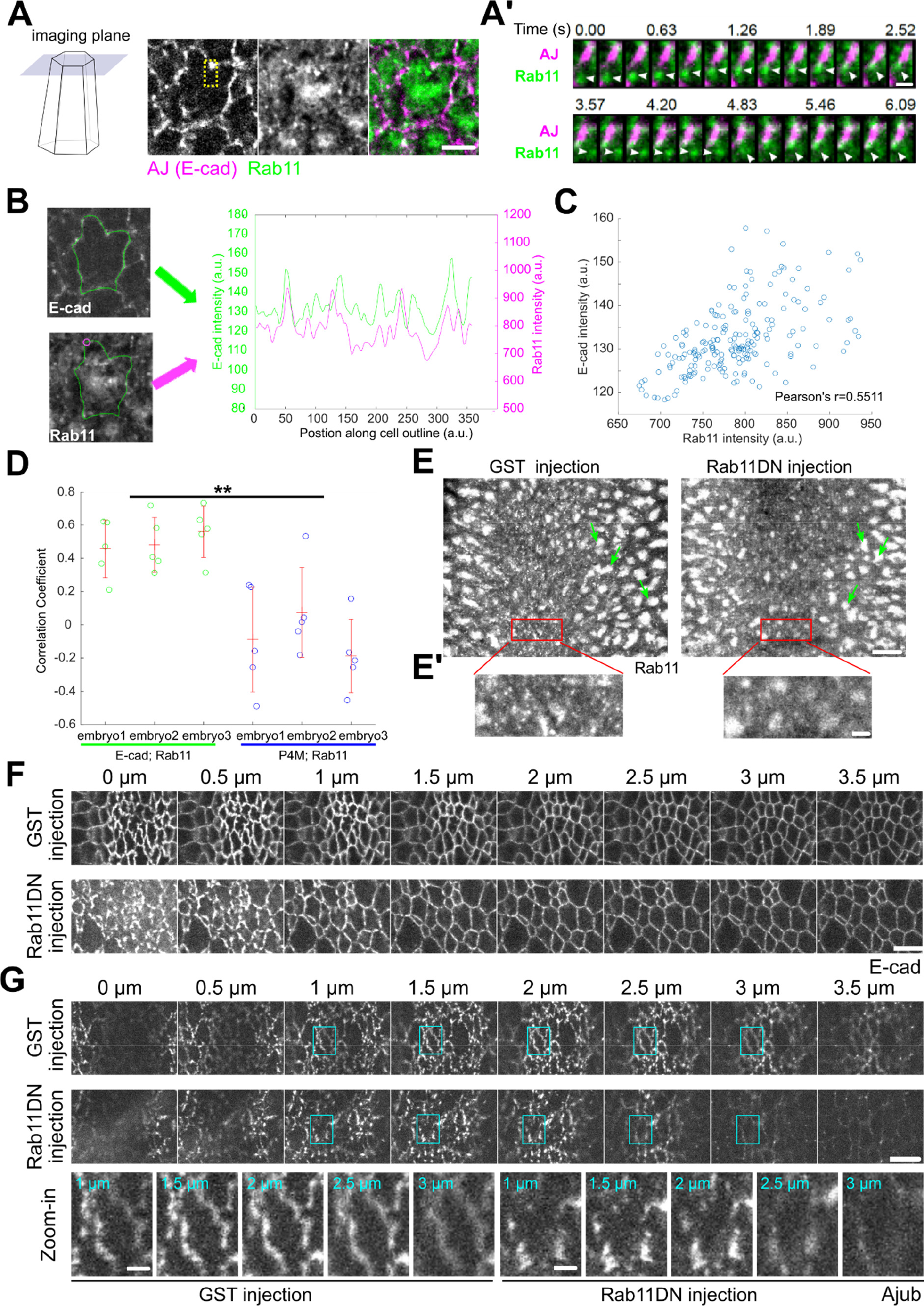
Rab11 reinforces the apical AJs during ventral furrow formation. **(A)** Capturing apical Rab11 vesicle dynamics with single-plane fast imaging. Example of a single constricting cell is shown. Scale bar, 5 μm. (A’) zoom-in view of (A) showing two Rab11 vesicles moving towards an adherens junction spot. Scale bar, 1 μm. **(B-D)** Analysis of spatial correlation between Rab11 vesicles and AJs. (B) The same constricting cell as shown in (A) is outlined in green. Magenta circle: an example sampling region within which Rab11 signal is measured. Right panel: the intensity profiles of E-cadherin and Rab11 along the indicated cell outline. (C) E-cadherin intensities for all points along the cell outline plotted against the corresponding Rab11 intensities. (D) Rab11 shows a significantly higher correlation with E-cadherin than with a general membrane marker P4M (N = 3 embryos for each genotype, 5 cells per embryo). Two tailed Student’s t-test was used for statistical comparison. **(E)** Injection of Rab11DN rapidly eliminates apical Rab11 vesicles but does not immediately affect the perinuclear Rab11 compartments (green arrows). Scale bar, 10 μm. (E’) Zoom-in view of (E). Scale bar, 2 μm. **(F)** Rab11DN injection affects the integrity of apical AJs in the constricting cells. 14 out of 19 Rab11DN injected embryos show fragmented apical AJs while 8 of 8 GST injected embryos show relatively continuous AJs. Scale bar, 5 μm. **(G)** Ajub shows reduced enrichment at the apical AJs during apical constriction in Rab11DN injected embryos compared to the control embryos (N = 4 embryos for each condition). Scale bar, 5 μm. Bottom, enlarged view of one cell (cyan boxes) as an example. Scale bar, 1 μm.

It remains unclear whether the targeting of Rab11 vesicles to the cell periphery is followed by fusion of vesicles with the plasma membrane, as demonstrated in some other systems (Grünfelder et al., 2003; Takahashi et al., 2012). We were not able to detect a clear, stable plasma membrane-associated signal for YFP-Rab11. Interestingly, a constitutively active form of Rab11 (Rab11CA), which is locked in its GTP-bound state, showed a prominent cell membrane localization in addition to the vesicle form (Figure S3A, B). As another evidence supporting the exocytic origin of the Rab11 vesicles, we found that injection of embryos with Brefeldin A (BFA), an Arf GEF inhibitor commonly used to block trafficking out of ER-Golgi system (Chardin and McCormick, 1999), resulted in rapid reduction of apical Rab11 vesicles and formation of large Rab11 aggregates between the cell apex and the nucleus (Figure S3C).

The enrichment of Rab11 vesicles near the apical AJs prompted us to ask whether Rab11 vesicles regulate the structure and function of AJs. To address this question, we inhibited Rab11 activity by injecting embryos with purified dominant negative Rab11 mutant proteins (Rab11DN), an approach previously used to reveal the function of Rab11 in cellularization (Pelissier et al., 2003). Rab11DN blocks the activation of endogenous Rab11 by binding to and sequestering the guanine nucleotide exchange factors (GEFs) for Rab11 (Ullrich et al., 1996). Importantly, the injection approach allowed stage-specific inhibition of Rab11, which was critical for bypassing the requirement for Rab11 in pre-gastrulation stages (Pelissier et al., 2003; Riggs et al., 2003; Tiwari and Roy, 2008). Injection of Rab11DN at mid/late cellularization completely prevented the accumulation of apical Rab11 vesicles during apical constriction (N = 5/5); whereas injection of Rab11DN close to the onset of apical constriction either eliminated apically accumulated Rab11 vesicles (in 4.4 ± 1.3 min, N = 24/31; Figure 8E; Video 6) or decreased their number (N = 7/31). These results suggest that active Rab11 is critical for both initiation and maintenance of apical accumulation of Rab11 vesicles. In contrast, the size and morphology of perinuclear Rab11 compartments were not immediately affected by injection (Figure 8E, green arrows). Injection of Rab11DN did not obviously affect the formation or morphology of apical Rab35 tubules, which have been previously implicated in facilitating endocytic membrane uptake from the apical domain during apical constriction (Jewett et al., 2017; Miao et al., 2019; Figure S4). Strikingly, injection of Rab11DN has a notable impact on the organization of apical AJs visualized by E-cadherin-GFP. In the control GST-injected embryos, apical AJs became more continuous and belt-like 6 minutes into apical constriction. In contrast, Rab11DN-injected embryos showed more fragmented apical AJs at comparable stage (Figure 8F). No difference was observed between the control and Rab11DN- injected embryos at the lateral region of the cells where E-cadherin is diffusely distributed (Figure 8F). Another apical AJ component, the Ajuba LIM protein Jub (Razzell et al., 2018), showed reduced enrichment at AJs upon Rab11DN injection (Figure 8G). Together, these results suggest that the apical Rab11 vesicles function to reinforce apical AJs during apical constriction.

### Rab11 regulates Myosin II organization and promotes apical constriction

Given the important role of AJs in the spatial organization of the supracellular actomyosin network (Martin et al., 2010; Sawyer et al., 2009), we examined the dynamics and organization of apical Myosin II in embryos injected with Rab11DN around the onset of gastrulation. Rab11DN injection at this stage did not cause any obvious defect in the apical activation of Myosin II (Figure 9A-C, GST vs. Rab11DN late injection). However, quantification of the rate of myosin flow towards the ventral midline showed a significantly lower rate in Rab11DN injected embryos compared to the control embryos (Figure 9D-G). This defect indicates that Rab11DN injection impaired apical constriction, which we confirmed by analysis of apical cell area changes (Figure S5A-E). The apical constriction defect was associated with frequent breaks within the apical Myosin II network as well as the underlying actin network (Figure 9H-J; Video 7). These myosin breaks occurred randomly across the constriction domain and were usually promptly reconnected, which prevented the complete tearing apart of the network. These observations suggest that the apical accumulation of Rab11 vesicles is important for maintaining the structural integrity of the apical actomyosin network. In support of this notion, similar myosin breaks have been observed in various conditions when apical Rab11 vesicle accumulation was abolished, such as disruption of microtubules (Ko et al.,2019), inhibition of dynein (Video 8) and BFA-mediated block of exocytosis (Figure S3C, asterisks; Video 9).

**Figure 9.**
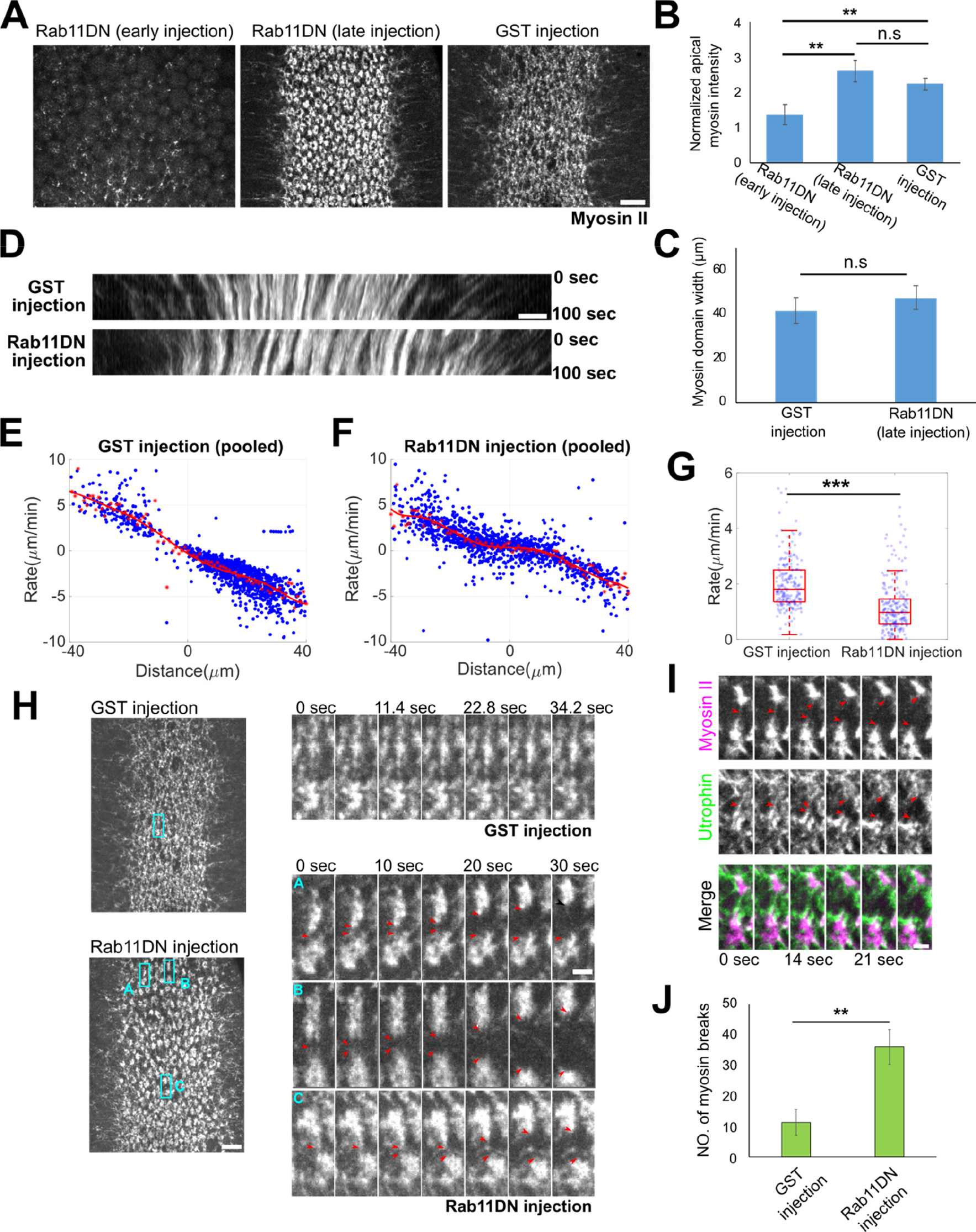
Acute inhibition of Rab11 results in defects in Myosin II organization and apical constriction. **(A-C)** Stage-dependent effect of Rab11DN injection on apical myosin network. (A): Injection of Rab11DN at early cellularization (> 30 min before onset of gastrulation) impairs apical myosin II activation (N = 3 embryos), whereas injection around the onset of gastrulation does not affect myosin II activation but causes defects in apical myosin organization (N = 4 embryos). GST injection: N = 3 embryos. Scale bar, 10 μm. (B, C): Quantification of apical myosin II intensity (B) and the width of myosin domain (C) at ∼6-7 min after the onset of gastrulation. Two tailed Student’s t-test was used for statistical comparison. **(D)** Example kymograph showing the movement of apical myosin II towards ventral midline at ∼6-7 minutes into gastrulation. Only the late Rab11DN injected embryos were analyzed in (D) – (K). Scale bar, 5 μm. **(E – G)** Quantification of apical constriction rate in embryos injected with GST and Rab11DN. (E, F) Rate of ventral movement of myosin structures as a function of their initial position relative to the ventral midline. Red asterisks: average velocity. Red curve: polynomial fit. (G) Rate for myosin structures located 10-15 μm away from the ventral midline. Two tailed Student’s t test was used for statistical comparison. **(H – J)** Rab11DN injection increases the frequency of myosin breaks during apical constriction. (H) Left, representative surface views of the supracellular myosin network. Scale bar, 10 μm. Right, zoom-in view showing myosin dynamics over time. Red arrowheads: myosin breaks. Scale bar, 2 μm. (I) Myosin breaks are associated with breaks in the underlying F-actin network (UtrophinABD-Venus). Scale bar, 2 μm. N = 3 embryos for each condition. (J) Quantification of the number of myosin breaks over a ∼100-sec time window at ∼6 – 7 min after apical constriction starts. Two tailed Student’s t test was used for statistical comparison.

Previous studies have demonstrated that apical constriction of the ventral mesodermal cells is pulsed, with successive constriction pulses interrupted by pauses when cells stabilize their constricted state (“ratcheting” mechanism; Martin et al., 2009; Mason et al., 2013). To ask whether Rab11DN injection affects pulsed apical constriction, we analyzed the dynamics of apical area change during ∼ 4 – 6 minutes after the onset of apical constriction (Methods). When Rab11 was inhibited, cells still exhibited pulsatile constriction behavior; there was no significant difference in the pulse interval between Rab11DN and control GST injected embryos (41.3 ± 17.8 s for GST injection, mean ± s.d., N = 21 cells from 2 embryos; 48 ± 20.4 s for Rab11DN injection, mean ± s.d., N = 26 cells from 2 embryos; Figure 10A-C).

**Figure 10.**
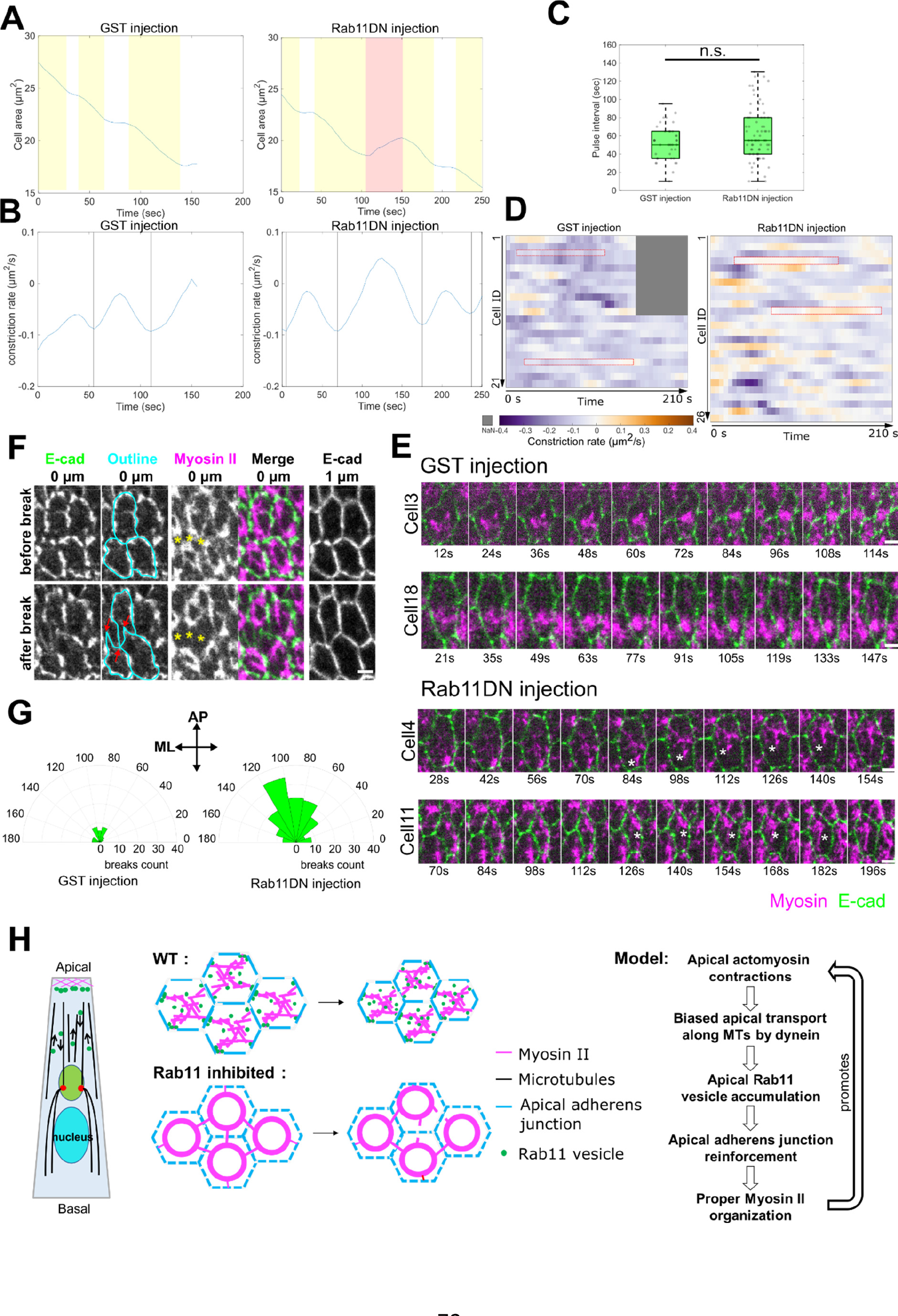
Rab11DN injection results in increased frequency of apical re-expansion during apical constriction. **(A, B)** Example cell area change and constriction rate over time in GST and Rab11DN injected embryos. Yellow shaded area, constriction phase; white area, stabilization phase; red shaded area, apical re-expansion. Vertical lines in B: peak of constriction rate. **(C)** Rab11DN injection does not affect the frequency of cell constriction pulses. Pulse interval was calculated as the interval between neighboring constriction peaks. **(D, E)** Rab11DN injected embryos exhibited increased frequency of apical re-expansion events. (D) Heatmap showing constriction rate of individual cell over ∼200 sec window during mid-apical constriction phase in GST and Rab11DN injected embryos. Positive constriction rate indicates apical re-expansion. N = 21 cells from 2 GST-injected embryos. N = 26 cells from 2 Rab11DN-injected embryos. Red boxes correspond to the examples shown in (E). Asterisks in (E): gaps resulted from myosin break event. Scale bars, 2 μm. (F) Apical AJ is pulled inward (red arrows) when myosin break (yellow asterisks) occurs in the adjacent cell. Scale bar, 2 μm. (G) Quantification of myosin break orientation in GST- and Rab11DN-injected embryos. Dataset from figure 9 (late injection) was used for this analysis. **(H)** Schematics of proposed actomyosin-dependent feedback mechanism.

However, Rab11DN injection resulted in an increased frequency of apical re-expansion after constriction pulses (Figure 10D), indicating that the cells were less capable of stabilizing their constricted state. Thus, the frequent myosin breaks caused by Rab11DN injection impaired the stabilization of the constricted state during pulsed constriction.

The apical re-expansion events in Rab11DN injected embryos were usually associated with visible local disconnection between the myosin structures and the cell-cell boundary (Figure 10E). During such events, apical AJs were often pulled to the opposite side from where myosin breaks happened (Figure 10F, red arrows). These observations suggest that the increased frequency of myosin breaks is due to impaired connections between junction and the actomyosin network, perhaps as a consequence of altered junction organization. In line with this view, we observed similar myosin breaks in embryos defective in *canoe*, the *Drosophila* Afadin homolog that regulates the linkage of the actin cytoskeleton to AJs during apical constriction (Video 10, Sawyer et al., 2009). Together, these results suggest that the apical Rab11 vesicles function to reinforce AJs and thereby strengthen the anchorage of the actomyosin network to the AJs. Interestingly, we found that myosin breaks and defect in apical constriction in the Rab11DN-injected embryos were more prominent during later phases of apical constriction (Video 7; Figure S5A-E), suggesting that the role of apical Rab11 vesicles becomes more important as tissue tension increases.

In addition to myosin breaks, we observed a second phenotype in the spatial organization of apical Myosin II. Instead of forming a supracellular meshwork with medioapically enriched myosin foci connected by myosin fibers, Myosin II formed ring-like circular structure at the apical domain of each constricting cell (Figure 9A). Interestingly, a recent study showed that loss of anisotropy in apical tension led to similar myosin ring formation in the constricting cells (Chanet et al., 2017). In Rab11DN-injected embryos, most myosin breaks predominantly occurred in the anterior-posterior (AP) orientation (Figure 10G), which is the direction in which tension is higher (Martin et al., 2010). These breaks may dissipate tension in the AP direction, thereby reducing tension anisotropy of the actomyosin network and causing myosin ring formation. In support of this view, the constricting cells in Rab11DN injected embryos exhibited no obvious increase of anisotropy in their apical geometry at the late stages of apical constriction, in sharp contrast to those in control embryos in which anisotropy continued to increase during apical constriction (Figure S5F). These observations led to an interesting hypothesis that the “myosin ring” phenotype in Rab11DN-injected embryos is a consequence of increased frequency of breaks in the supracellular actomyosin network.

### Distinct early and late functions of Rab11 in regulating apical Myosin II in mesoderm primordium

The lack of defects in apical myosin activation upon Rab11 inhibition apparently contradicted with previous findings that Rab11 is required for apical myosin activation in several other apical constriction-mediated tissue folding processes (Le and Chung, 2021; Ossipova et al., 2014, 2015). In these studies, Rab11 was typically inhibited at a stage much earlier than the onset of apical constriction. We found that injection of Rab11DN proteins during early cellularization (> 30min before the onset of gastrulation) indeed resulted in a strong reduction or lack of apical myosin accumulation during gastrulation (Figure 9A, B). Therefore, through acute inhibition of Rab11, we were able to resolve the early and late functions of Rab11 in apical constriction and identify a previously unappreciated function of Rab11 in the feedback regulation of apical actomyosin network post Myosin II activation (Figure 10H).

## Discussion

Our work reveals an intimate interplay between actomyosin network and Rab11-mediated vesicular trafficking during apical constriction mediated epithelial folding. Combining live imaging with genetic, optogenetic and pharmacological approaches, we show that during ventral furrow formation, apical actomyosin induces dynein- and microtubule-dependent, biased transport of Rab11 vesicles towards the cell apices, which facilitates the enrichment of the vesicles at apical AJs. By acute elimination of apical Rab11 vesicles through Rab11DN injection, we further present evidence that these vesicles are important for maintaining the structural integrity of the AJs and for preventing undesired breaks between actomyosin network and the AJs during apical constriction. These results suggest an actomyosin-dependent, Rab11-mediated feedback mechanism that serves to reinforce the connections between the actomyosin network and apical AJs (Figure 10H). In further support of the proposed mechanism, we found that acute disruption of Rab11 vesicle transport or blocking trafficking through the Golgi apparatus resulted in similar defects in the integrity of the supracellular actomyosin network. Finally, we show that this feedback mechanism is separate from the function of Rab11 in promoting apical Myosin II activation as previously reported in other systems. The mechanism identified in this work allows the tissue to promptly adapt to the increased tissue tension during apical constriction and maintain the integrity of the force generation machinery in the process of tissue folding.

The apical enrichment of Rab11 vesicles is closely associated with the formation of intact actomyosin network at the apical domain of the cell, as evidenced by the strict correlation between the two processes in a variety of conditions we tested. In particular, we show that accumulation of myosin at the apical cortex without F-actin (and thereby lacking contractility) is not sufficient to induce vesicle accumulation, and that the directional bias in Rab11 vesicle transport diminishes immediately after acute actomyosin inactivation. The outcomes of these tests also let us rule out the requirement of actomyosin-AJ coupling and apical shrinking for Rab11 vesicle accumulation. Since actomyosin networks are intrinsically contractile, our findings raise an intriguing possibility that the biased Rab11 vesicle transport is subjected to mechanosensitive regulations. In recent years, it has been increasingly appreciated that cell and tissue-scale mechanical forces can impact intracellular membrane trafficking. For example, high membrane tension can suppress the formation of endocytic vesicles (Wu et al., 2017) and promote vesicle fusion with lipid bilayer (Shillcock and Lipowsky, 2005; Staykova et al., 2011). Mechanical stresses can also impact vesicle transport and distribution in neurons (Ahmed and Saif, 2014; Ahmed et al., 2012; Siechen et al., 2009) and trigger regulated exocytosis in various contexts (Boycott et al., 2013; Gauthier et al., 2011; Khandelwal et al., 2013). The rapid alteration in Rab11 vesicle transport in response to actomyosin activities provides a potential example of mechanical regulation of intracellular trafficking in the context of tissue morphogenesis.

The way how apical constriction affect Rab11 vesicle transport is unknown. Apical constriction may cause changes in the activity of the motor complexes that drive the motion of Rab11 vesicles. Regulation of dynein-mediated transport can occur at multiple levels, including Rab11-adaptor binding, adaptor activities, adaptor-motor binding, dynein recruitment to microtubules, and motor activity itself (Dillman and Pfister, 1994; Fu and Holzbaur, 2013; Horgan et al., 2010; Moughamian et al., 2013; Otani et al., 2011; Vaughan et al., 2002; Wang et al., 2019). Alternatively, apical actomyosin contractions may bias the direction of transport by generating a “sink” that retains the vesicles at the apical side, thereby inhibiting the initiation of the reverse, basally directed transport of the vesicles.

### Future research is required to further tease apart these possibilities

Another important question remained to be address is the molecular basis of the myosin breaks observed in Rab11DN injected embryos. Rab11 may regulate apical constriction by regulating endocytic recycling of Fog and GPCR (Fuse et al., 2013; Jha et al., 2018; Pouille et al., 2009). However, alteration in Fog-GPCR signaling would be expected to affect apical myosin II activation, which was not observed in our experiments upon acute Rab11DN injection. Rab11 may also be involved in Rab35-mediated endocytosis of apical membrane, which has been shown to facilitate ratcheted constriction (Jewett et al., 2017; Miao et al., 2019). However, we found that the apical Rab11 vesicles are not derived from endocytosis and that acute injection of Rab11DN did not obviously affect the Rab35 compartments.

Instead, we show that in Rab11DN injected embryos, the appearance of frequent myosin breaks was associated with severe fragmentation of apical AJs, raising the possibility that the function of Rab11 in maintaining the integrity of the actomyosin network is attributed to its role in regulating apical AJs. Rab11 recycling endosomes have been shown to regulate the delivery of E-cadherin to the AJs in a number of other cell and tissue contexts (Langevin et al., 2005; Le Droguen et al., 2015; Lock and Stow, 2005; Woichansky et al., 2016; Yashiro et al., 2014). However, we did not detect E-cadherin-GFP signal on Rab11 vesicles during apical constriction. It is possible that the signal was too weak to be detected by the method we used. Alternatively, these vesicles may function to transport other junction components or regulators. Finally, Rab11 vesicles may regulate the connection between AJ and actomyosin network by delivering actin regulators to AJs, as dynamic actin turnover at AJs has been shown to impact the detachment and reattachment of actomyosin network to AJs (Jodoin et al., 2015). Future studies identifying the cargos of the Rab11 vesicles during apical constriction will be the key to understand the molecular function of these vesicles.

## Materials and methods

### Fly stocks and genetics

*Drosophila melanogaster* flies were grown and maintained at 18°C and crosses were raised and maintained at room temperature (21 – 23°C) unless otherwise mentioned. All flies were raised on standard fly food. For embryo collection, flies with corresponding genotype were used to set up cages and maintained at 18°C, and embryos were collected from apple juice agar plate containing fresh yeast paste.

For most experiments, embryos expressing endogenously tagged YFP::Rab11 (Dunst et al., 2015) (BDSC Stock#62549) were imaged to visualize Rab11 vesicles. We also generated UAS-mCherry::Rab11 transgenic flies for dual-color imaging with other GFP-tagged markers (Figure 6A and B, Figure 8A, Figure S2C and Figure S4). For lines with UAS-driven transgenes or shRNA, we used GAL4 driver lines carrying matα4-GAL-VP16 (denoted as “mat67” on the 2nd chromosome and “mat15” on the 3rd chromosome) to drive maternal expression in the embryo through either direct cross or recombination.

The following fly lines were generated for dual-color imaging of Rab11 with other markers in this study:

– Sqh::mCherry; YFP::Rab11

– Gap43::mCherry/Cyo; YFP::Rab11

– mat67 mCherry::Rab11; mat15 GFP::Jupiter

– mat67 mCherry::Rab11; mat15 E-cadherin::GFP

– mat67 mCherry::Rab11;mat15 mCherry::P4M

Figure S1: To examine other organelle behavior during apical constriction, the following lines were either directly used or first crossed to mat67; mat15 flies in order to obtain embryos expressing the corresponding fluorescent marker:

– UAS-Arf79F::GFP (BDSC Stock#65850)

– UAS-KDEL::GFP (BDSC Stock#9898)

– UAS-YFP::Rab7 (BDSC Stock#23641)

– YFP::Rab5 (BDSC Stock#62543)

– YFP::Rab8 (BDSC Stock#62546)

Figure 2C: To knock down Myosin II, female flies of a TRiP *zipper*; YFP::Rab11 stock generated from a TRiP *zipper* RNAi line (BDSC Stock#37480) were crossed to male flies from stock mat67 Sqh::mCherry; mat15 YFP::Rab11 to obtain F1 TRiP *zipper*/mat67 Sqh::mCherry; YFP::Rab11/mat15 YFP::Rab11 flies, and embryos from these F1 flies were collected for imaging.

Figure 2D: To generate ventralized embryos, *Spn27A1*/Cyo; Sqh::mCherry YFP::Rab11/TM3 was crossed to Df(2L)BSC7/Cyo; Sqh::mCherry YFP::Rab11/TM3 to obtain *Spn27A1*/ Df(2L)BSC7; Sqh::mCherry YFP::Rab11 flies, and embryos from these flies were collected for imaging. *Spn27A* encodes a serine protease inhibitor that regulates dorsal-ventral patterning in early embryos by inhibiting the Toll-Dorsal pathway (Ligoxygakis et al., 2003). Consistent with previous reports, *Spn27A* mutant embryos showed a ventralized phenotype with expanded apical Myosin II activation domain during gastrulation (Figure 2D; Ligoxygakis et al., 2003).

Figure 3C: To knock down α-catenin, female flies of TRiP *α-catenin* RNAi line (BDSC Stock# 33430) were crossed to male flies from stock mat67 Sqh::mCherry; mat15 YFP::Rab11 to obtain F1 +/mat67 Sqh::mCherry; TRiP *α-catenin* /mat15 YFP::Rab11 flies, and embryos from these F1 flies were collected for imaging. Consistent with previous reports, disrupting AJs did not prevent apical myosin activation, but rather impaired the connection between the actomyosin network and the cell-cell boundaries. As a result, Myosin II coalesced into concentrated spots at the center of each individual medioapical domain (Martin et al., 2010).

Figure 4A-C: To overexpress Fog, female flies from UAS-Fog stock were crossed to male flies from stock mat67 Sqh::mCherry; mat15 E-cad::GFP or mat67 Sqh::mCherry; YFP::Rab11 to obtain corresponding F1 flies, and embryos from these F1 flies were collected for imaging. As shown in a previous study, ectopic myosin activation in dorsal cells by ectopic expression of Fog does not cause cell apex shrinking or tissue indentation, thereby providing a scenario where actomyosin contractility and apical cell area reduction is decoupled (Morize et al., 1998).

Figure 7A-F: Female flies from stock CRY2::mCherry::Rho1DN YFP::Rab11/TM6C; CIBNpm/FM7 were crossed to male flies from stock mat67 Sqh::mCherry; mat15 YFP::Rab11 to generate mat67 Sqh::mCherry/+; CRY2::mCherry::Rho1DN YFP::Rab11/mat15 YFP::Rab11; CIBNpm/+ flies, and embryos from these flies were collected for imaging.

Figure 7G, H: Female flies from stock CRY2::mCherry::Rho1DN /TM6C; CIBNpm/FM7 were crossed to male flies from stock mat67 Sqh::mCherry; mat15 GFP::Jupiter to generate mat67 Sqh::mCherry/+; CRY2::mCherry::Rho1DN / mat15 GFP::Jupiter; CIBNpm/+ flies, and embryos from these flies were collected for imaging.

Figure 8, 10, S5: Sqh::mCherry E-cadherin::GFP was used for Rab11DN injection experiment for examining adherens junction phenotype and quantifying apical area change. Sqh::mCherry mat67; mat15 UAS-Jub::GFP was used for figure 8G.

Figure 9A-H: embryos from SqhmCherry; YFP-Rab11 flies were used.

Figure 9I: embryos from mat67 Sqh::mCherry; mat15 UtrABD::GFP were used in Rab11DN injection experiment for examining apical actin network.

Figure S2C: embryos from mat67 mCherry::Rab11/YFP::Rab5 flies were used.

Figure S2D, E: The fly stock used for temperature shift experiment is *shi^ts^*/ *shi^ts^*; Sqh::mCherry; YFP::Rab11.

Figure S3A, B: To visualize the localization of constitutively active Rab11, UAS- YFP::Rab11Q70L (BDSC Stock#9791) was crossed to mat67; mat15 males, and embryos from F1 females were used for imaging.

Figure S4: embryos from mat67 mCherry::Rab11/UAS-YFP::Rab35 were used.

Video 10: A TRiP *canoe* RNAi line (BDSC Stock#38194) was used for *canoe* knockdown. Female flies from the TRiP *canoe* stock were crossed to male flies from stock mat67 Sqh::mCherry; mat15 E-cad::GFP to obtain F1 TRiP *canoe*/mat67 Sqh::mCherry; +/mat15 E- cad::GFP flies, and embryos from these F1 flies were collected for imaging.

### Molecular cloning and generation of transgenic fly lines

To make construct for in vitro expression of recombinant dominant negative Rab11 (S25N) protein, dominant negative Rab11 coding sequence was PCR amplified from genomic DNA of UAS-YFPRab11DN (BDSC Stock#9792) and inserted into pGEX-6p-1 vector (a gift from the Griffin lab, Dartmouth College) using BamHI and XhoI restriction sites.

To make constructs for transgenic fly lines, mCherry-P4M double strand DNA was synthesized in vitro (Integrated DNA Technologies) and inserted into a fly transformation vector (pTiger, courtesy of S. Ferguson, State University of New York at Fredonia, Fredonia, NY, USA) using NotI and NheI restriction site to generate the pTiger-mCherry-P4M plasmid.

To make pTiger-mCherry-Rab11 construct, wild type Rab11 coding sequence was PCR amplified from genomic DNA of UAS-YFP::Rab11 (BDSC Stock# 9790) and cloned into pTiger-mCherry-P4M construct using SpeI and NheI restriction sites to replace the P4M sequence.

The resulting pTiger-mCherry-P4M and pTiger-mCherry-Rab11 constructs were sent to BestGene, Inc., for integration into either attP2 or attP40 site using the phiC31 integrase system (Groth et al., 2004).

### Live imaging and optogenetics

Embryos were dechorionated in 3% bleach, rinsed with water 12 times and mounted in water in a 35 mm MatTek glass-bottom dish (MatTek Corporation). Unless otherwise mentioned, all images were obtained using a Nikon inverted spinning disk confocal microscope equipped with the perfect focus system and Andor W1 dual camera, dual spinning disk module. An CFI Plan Apo Lambda 60×/1.40 WD 0.13 mm Oil Objective Lens objective was used for imaging at room temperature. YFP and GFP tagged proteins were imaged with a 488-nm laser and mCherry tagged proteins were imaged with a 561-nm laser. Images in Figure 1A, S1A and B were obtained using an upright Olympus FV-MPERS multiphoton microscope equipped with the InSight Deep Laser System with an Olympus 25X/1.05 water dipping objective (XLPLN25×WMP2). Images in Figure 1D, 2A, 2D, S2, S3 were obtained using a Zeiss Axio Observer laser scanning confocal microscope (LSM 880).

For optogenetic experiments, flies were kept in the dark and live sample preparation was performed under red light. Embryos were first imaged with 561-nm laser to visualize Myosin II signal to determine the developmental stage of the embryo. Once the embryo reached the desired stage, two-color imaging with both 488-nm and 561-nm lasers were carried out, where the 488-nm laser was used for both optogenetic stimulation and YFP-Rab11 visualization. The Opto-Rho1DN optogenetic tool used in this experiment is based on the CIBN-CRY2 system (Guglielmi et al., 2015; Kennedy et al., 2010). Blue-light dependent recruitment of a CRY2-Rho1DN fusion protein to plasma membrane-anchored CIBN blocks activation of endogenous Rho1, causing rapid dissociation of Myosin II and F-actin from the cell apices, thereby acutely inhibits apical actomyosin contractility (Guo et al., 2021).

For the temperature shift experiment with *shi^ts^* mutants, a Zeiss Axio Observer laser scanning confocal microscope (LSM 880) with an incubation chamber was used. Embryos subjected to restrictive temperature were prepared at room temperature and then transferred to an incubation chamber, which was preheated to 32°C. A 40X/1.3 numerical aperture oil-immersion objective, and 488-nm argon laser and 561-nm laser were used for imaging.

### Rab11 dominant negative protein expression and purification

pGEX-6p-1-Rab11DN plasmid was transformed into E.coli (BL21(DE3), New England BioLabs). An empty vector with only GST coding sequence was also transformed as control. After IPTG induction, bacteria were resuspended in 32 mL lysis buffer (50 mM Tris pH7.5, 150 mM NaCl, 0.1% Triton X-100, 1 mM PMSF, 1 mM DTT) and lysed with 0.25 mg/mL lysozyme incubation on ice for 2 h followed by sonication (6 rounds of 10 pulses every 1 minute). GST and GST-Rab11DN were purified from the supernatant with Glutathione Sepharose 4B GST-tagged protein purification resin (GE17-0756-01, Sigma-Aldrich) using batch method. Eluted protein was dialyzed using slide-A-Lyzer dialysis cassettes (Cat#66380, Thermo Scientific) with 50 mM Tris buffer (pH8.0) and further concentrated to a final concentration of ∼200 μM with Amicon Ultra-0.5mL centrifugal filters (Cat#UFC501024, Millipore Sigma).

### On-stage drug / protein injection

Embryos at cellularization stage were prepared as previously described, then mounted ventral side down on a 50 x 25 mm glass coverslip pre-covered with a thin layer of glue. Embryos were dried for 10-15 minutes in a desiccator. Embryos were then covered with halocarbon oil (halocarbon 700/halocarbon 27 = 3:1). A homemade injection device mounted next to the spinning disk confocal microscope was used for on-stage injection. Embryos were injected laterally into the ventral part of the embryo. 50 mM Y-27632 (Enzo Life Sciences), 25 mg/mL colchicine (Sigma-Aldrich), 5 mg/mL Cytochalasin D (Sigma-Aldrich), 0.5 μg/μL FM4-64 (Invitrogen^TM^) and 10 mg/mL Brefeldin A (Sigma-Aldrich) were used for corresponding experiment. For Rab11DN injection, Triton X-100 was added to a final concentration of 0.1% to prevent the solution from clogging the needle. For dynein and GST antibody injection, commercial monoclonal antibodies against cytoplasmic dynein intermediate chain (Cat#sc-13524, Santa Cruz Biotechnology) and monoclonal antibody against GST (Cat#sc-138, Santa Cruz Biotechnology) were first concentrated to 2 mg/mL with Amicon Ultra-0.5 mL centrifugal filters before injection (Cat#UFC501024, Millipore Sigma). For FM4-64 injection, the membrane dye was injected into the perivitelline space at late cellularization to label the cell membrane and monitor the localization of endocytosed membrane dye. For all other injections, the indicated reagents were injected into the embryo.

### Image processing and analysis

All images were processed using ImageJ (NIH) and MATLAB (MathWorks). For the following figures, due to laser power fluctuation and the extent of photobleaching with different imaging duration, the contrast was adjusted to make the cytoplasmic signal comparable: Figure 2A, 2B (Rab11), 2C (Myosin II), 2D (Myosin II and Rab11), S3(Myosin II and Rab11), 6C (Rab11), 6H (Myosin II and Rab11), 8E, 8F, 9A.

For 3D segmentation and rendering of Rab11-positive structures (Figure 1F-H), we first segmented a group of presumptive mesodermal cells in membrane channel using Embryo Development Geometry Explorer (EDGE), a MATLAB-based software (Gelbart et al., 2012) to generate individual cell mask. Then we used ilastik, a machine learning based software (Berg et al., 2019) to segment Rab11 vesicles and perinuclear compartments in the Rab11 channel (Pixel classification) to generate a probability map. A custom MATLAB script was used to convert the probability map for vesicle and perinuclear compartment to binary mask. These masks were applied to original image to remove background signal and 3D rendering of resulting images was created using Nikon Elements software. To generate 3D view of Rab11 structures in a single cell, the cell mask generated from EDGE was applied to the processed Rab11 images before 3D rendering. In Figure 1F and 1G, the original images are generated by maximum intensity projection rendering method while the rest of images are generated with alpha blending method.

For vesicle tracking in embryos expressing YFP-Rab11 and mCherry-Sqh (Figure 5A-E, Figure 7A-E), a 13.8 μm x 6.9 μm region of interest including approximately one constricting cell at 50% egg length was selected, and Rab11 vesicles were manually marked frame by frame for each trajectory using a multi-point tool in ImageJ. The x-y coordinates of individual vesicles over time were then exported. A MATLAB script was written to reconstruct the trajectories based on the coordinate information. The direction of each trajectory was defined by the relative translocation of the vesicle along apical basal axis between trackable start and end positions. All trajectories were grouped into two categories based on their direction (apical or basal) for further analysis of parameters, including trajectory count, average velocity, trajectory length and trajectory duration.

To validate the vesicle tracking analysis (Figure 5F, G), we performed the following analyses. First, we imaged embryos expressing both myosin and membrane markers at multiple z- planes. This approach allowed us to map the position of the single plane image to the cross- section view of the embryo (Figure 5F). The results confirmed that our single-plane imaging approach captured the cells located at the medial-lateral boundaries of the constricting domain. We noticed that although all of these cells showed apical myosin accumulation, some cells were constricted apically, whereas others appeared to be in the process of apical constriction or even stretched by the more centrally localized constricting cells. These observations were consistent with a recent report demonstrating that apical myosin contractility is graded from the center of the constriction domain (Denk-Lobnig et al., 2021). Second, we imaged embryos expressing both a membrane marker and mCherry-Rab11 (overexpression) and quantified transport events only in cells displaying obvious apical shrinking. We found no significant difference between the results from the original and the new analyses (Figure 5G). These additional analyses also indicate that apical actomyosin accumulation, but not apical area reduction, is the determining factor for the biased transport of Rab11 vesicles, consistent with the observation in dorsal cells with ectopic Fog expression (Figure 4A-C).

For quantification of vesicle density in 2D (Figure 4D-F), 3 separate ROIs covering most of the constriction region were selected, with each ROI enclosing a relatively flat piece of tissue surface (based on the apical Myosin II signal). For each ROI, a Z position near the apical surface where most vesicles were accumulated was selected for quantification. Vesicles were detected using the Find Maxima function in ImageJ in combination with manual correction to ensure accuracy of the counting. For vesicle counting in 3D (Figure 4D-F), the FIJI plugin 3D Maxima Finder was used and the noise tolerance parameter was determined based on average intensity of the entire image stack (Ollion et al., 2013).

The colocalization between Rab11-mCherry and E-cadherin-GFP (Figure 8B-D) was quantified in embryos at the stage of early ventral furrow formation. For each embryo, 5 non-neighboring constricting cells were manually segmented using a multi-point tool to mark the vertices of the cell outline. A MATLAB script was written to import the coordinates of these vertices to create the cell outline. For each cell outline, a series of closely spaced and evenly distributed sampling points were generated along the outline. For each sampling point, E-cadherin-GFP signal intensity on that single pixel was extracted as the junction signal, and average Rab11-mCherry signal intensity within a radius of 5 pixels (∼0.54 μm) was calculated as Rab11 signal for the same point. For each cell, a scatter plot of E-cadherin intensity over Rab11 intensity for all the sampling points was generated, and a correlation coefficient was calculated. A similar analysis was performed between P4M-GFP and Rab11- mCherry as a control.

For Myosin II intensity quantification (Figure 9B), 100-pixel by 100-pixel ROIs (10.8 μm X 10.8 μm) were taken from the apical Myosin II domain. Average intensity R was measured inside the ROI on that plane, and background noise at the same plane (Rbg) was measured as the mean intensity of a random ROI outside the embryo. To assess cytoplasmic myosin intensity (C), we used the same ROI position but at a slightly deeper Z below the apical myosin network, the background noise at that plane (Cbg) was measured as described above. The normalized Myosin II intensity I = (R-Rbg) / (C-Cbg). Embryos at ∼6 – 7 min after the onset of gastrulation were used for quantification.

To quantify the rate of apical constriction based on the movement of apical Myosin II structures (Figure 9D-G), kymographs were generated from the same dataset of the above-mentioned 100-sec myosin movies. For each embryo, a series of myosin kymographs were generated by average projection over every 30 pixels (3.2 μm) along anterior posterior axis. Bandpass filter in ImageJ were used to remove noise. A custom MATLAB script was used for automatic segmentation of individual myosin traces and fitting with lines. Rate of Myosin II moving towards ventral midline was extracted by calculating the slope of each myosin trace. Velocity of Myosin II at ventral midline was subtracted to correct for the drift of ventral midline itself. The rate of myosin movement was then plotted against the distance of the corresponding myosin structure to ventral midline at the time when tracing started.

For quantification of myosin breaks (Figure 9J, 10G), a maximum projection over 3 – 5 μm thick Z stacks that include all apical Myosin II signal was generated. A ∼100-sec time window starting from the time point when the Myosin II domain at the ventral surface exhibits a relatively clear domain boundary (∼6 – 7 min after the onset of gastrulation) was selected to count the number of myosin break events. At the site where myosin break occurs, the myosin fiber usually snaps and the two ends rapidly retract, which creates a clear gap between neighboring myosin clusters.

To analyze the cell area dynamic from fast movies (Figure 10A-D), the time lapse movie of a subapical single slice from E-cad channel was first rearranged to convert T dimension into Z dimension. Individual cells in the resulting image were segmented in ilastik using carving method (Berg et al., 2019). Once segmented, cell area, constriction rate and pulse duration were calculated using a customized MATLAB script. In order to find the peaks for the constriction rate, the constriction rate curve was first smoothed using MATLAB smooth function with a span of 5. The rate was then inverted to convert valleys into peaks and MATLAB findpeaks function was used to locate the peaks and pulse duration was calculated as the interval between neighboring peaks.

To quantify the rate of apical constriction from E-cadherin-GFP movies (Figure S5A, B), images at 6 – 7 μm below the apical surface were used to measure the cell area change over time. A group of constricting cells at the ventral most region of the embryo were outlined frame by frame using multi-point tool in ImageJ. The area of the same cell group over time was then calculated and plotted using MATLAB.

To analyze the distribution of cell area in the same embryo at 2 min, 5 min and 8 min after onset of gastrulation (Figure S5C-E), individual constricting cell was segmented using Embryo Development Geometry Explorer (EDGE), a MATLAB-based software (Gelbart et al., 2012). Individual cell area was then calculated, and the distribution histogram of cell area was plotted. Cell anisotropy was calculated based on the same segmented cell dataset (Figure S5F). Two orthogonal (anteroposterior and mediolateral) lines passing the centroid of the cell were drawn across the segmented cell, and the cell anisotropy was calculated as the ratio between the anteroposterior intercept by the cell boundary and the mediolateral intercept.

## Statistics

Statistical comparisons were performed using two tailed one-sample t test or two-tailed Student’s t tests. Sample sizes can be found in figure legends. *p* values were calculated using MATLAB ttest (Two tailed one-sample t test) and ttest2 function (Two tailed Student’s t test).

## Supplemental Material

Figure S1 shows changes in 3D Rab11 structures during ventral furrow formation as well as a comparison to other endomembrane compartments. Figure S2 shows that Rab11 vesicles are not derived from endocytosis. Figure S3 shows the localization of constitutively active Rab11 during ventral furrow formation and the impact of BFA injection on Rab11 and Myosin II. Figure S4 shows that Rab11DN injection does not obviously affect Rab35 structures. Figure S5 shows that acute inhibition of Rab11 results in reduced rate of apical constriction. Video 1 shows myosin network relaxation immediately after cytoD injection. Video 2 shows Rab5, Rab11 localization relative to endocytosed vesicles after FM4-64 membrane dye injection.

Video 3 shows the impact of acute disruption of MTs on Rab11 vesicle transport. Video 4 shows the impact of dynein inhibition on Rab11 vesicle transport. Video 5 shows the dynamics of Rab11 vesicles near apical adherens junctions. Video 6 shows the rapid disappearance of Rab11 vesicles after Rab11DN injection. Video 7 shows that injection of Rab11DN causes frequent breaks within the apical actomyosin network. Video 8 shows that injection of dynein antibody affects apical Rab11 vesicle accumulation and causes frequent myosin breaks. Video 9 shows that injection of BFA results in abnormal Rab11 structures and causes frequent myosin breaks. Video 10 shows that knockdown of Canoe causes frequent myosin breaks.

## Acknowledgments

We thank members of the He lab and the Griffin lab for discussions throughout this work; Yashi Ahmed, Charles K.Barlowe and Adam Martin for valuable comments on the manuscript; Ann Lavanway and Zednek Svindrych for technical support with imaging; Robert Robertson for help with on-stage injection setup. We thank the Wieschaus lab and the De Renzis lab for sharing reagents. We thank the Bloomington *Drosophila* Stock Center for fly stocks. This study was supported by NIGMS ESI-MIRA R35GM128745 and American Cancer Society Institutional Research Grant #IRG –82-003-33 to B.H. The study used core services supported by STANTO15R0 (CFF RDP), P30-DK117469 (NIDDK P30/DartCF), and P20-GM113132 (bioMT COBRE).

## Author Contributions

W.C. and B.H. designed the study. W.C. performed the experiments and analyzed the data.

W.C. and B.H. wrote the manuscript.

---

**Figure S1.**
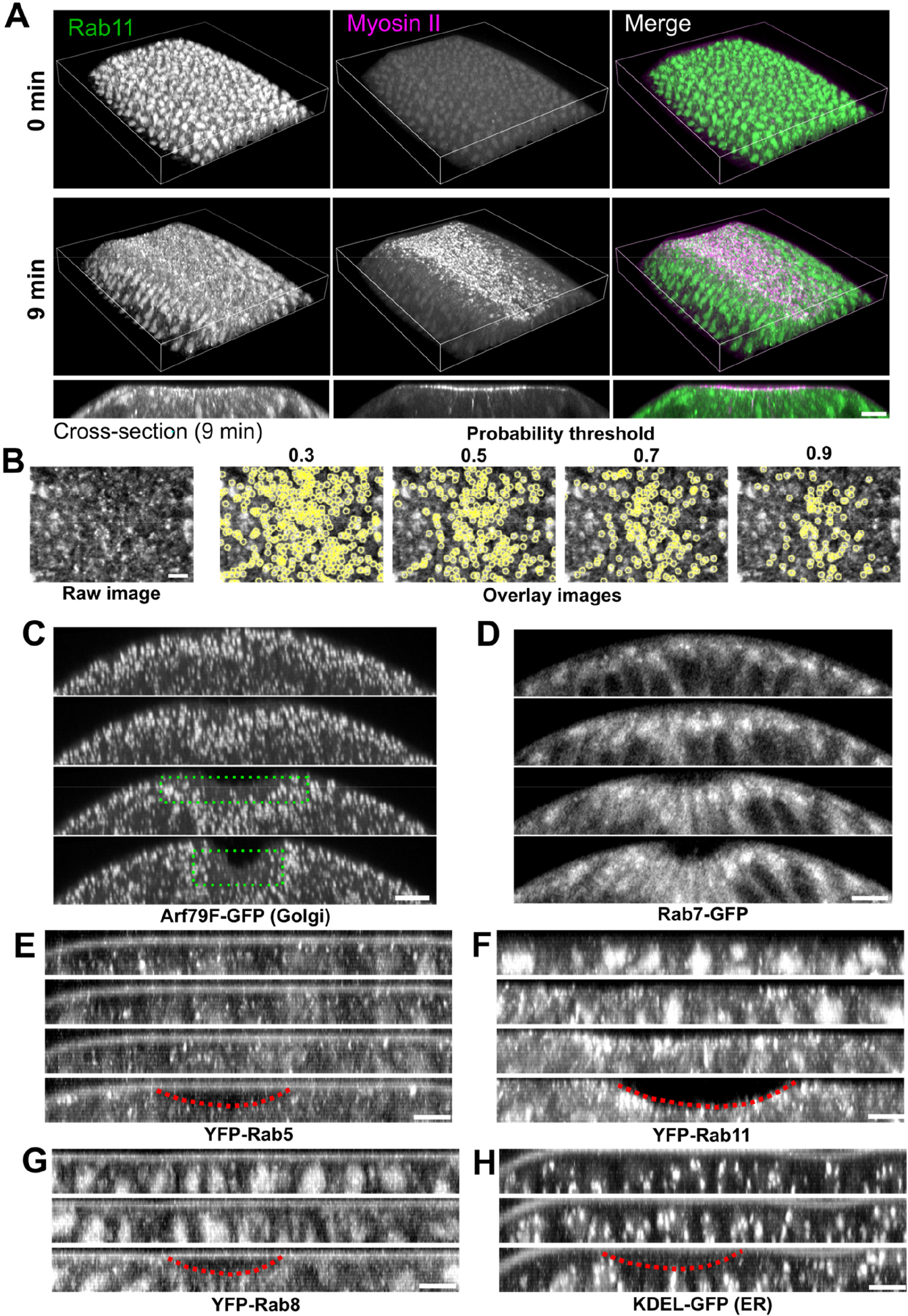
Localization of Rab11 and other endomembrane compartments during ventral furrow formation. **(A)** 3D reconstruction of the ventral part of an embryo expressing YFP-Rab11 and mCherry- Sqh during ventral furrow formation showing apical accumulation of Rab11 vesicles in constricting cells. Magenta box indicates the apical region of the constricting cells. Bottom panel are the cross-section view images of the same embryo. Scale bar, 10 μm. **(B)** Vesicle segmentation using different probability threshold. Vesicles are segmented based on the probability map generated by ilastik. Threshold of 0.7 gives best quality of segmentation. **(C-H)** Localization change of different compartment markers in the ventral cross-section view of embryos. from late cellularization (top image in each panel) to early gastrulation. (C) Golgi marker Arf79F. Green dotted box shows the lack of Golgi puncta in the apical region of the constricting domain. (D) Late endosome marker Rab7. (E) Early endosome marker YFP-Rab5. (F) Recycling endosome marker YFP-Rab11. (G) YFP-Rab8. (H) ER marker KDEL-GFP. Among all the organelles examined here, Rab11 positive structures exhibit the most striking morphology and localization changes during ventral furrow formation. Images in A and B were taken on a multiphoton microscope. Scale bars, 10 μm. Images in C – F were taken on a spinning disk confocal microscope. Red dotted lines indicate the ventral furrow. Scale bars, 5 μm. The cross-section views were made by maximum intensity projection over 4 – 5 μm.

**Figure S2.**
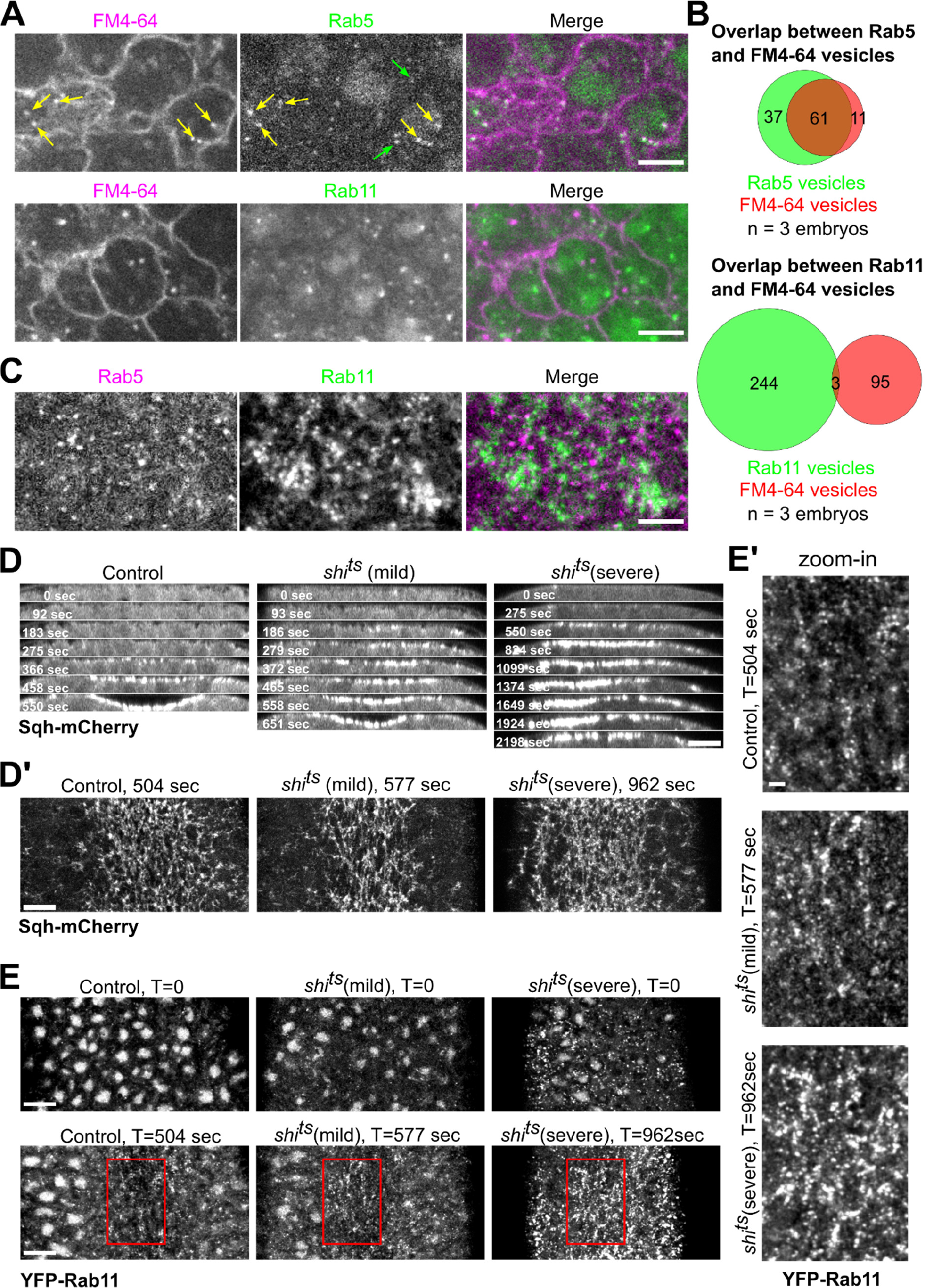
Apical Rab11 vesicles are not derived from endocytosis. **(A)** Endocytosed positive vesicles labeled by membrane dye FM4-64 extensively colocalize with YFP-Rab5 vesicles (85%, 72 vesicles from 3 embryos) but not YFP-Rab11 vesicles (3%, 98 vesicles from 3 embryos) during apical constriction. Green arrows mark Rab5 vesicles that do not colocalize with FM4-64 vesicles and yellow arrows mark Rab5 vesicles that colocalize with FM4-64 vesicles. Scale bar, 5 μm. **(B)** Quantification of colocalization between endocytosed FM4-64 vesicles and corresponding Rab vesicles. **(C)** YFP-Rab5 and mCherry-Rab11 vesicles do not show colocalization. Scale bar, 5 μm. **(D, D’)** Top panels (D): Montage showing Sqh-mCherry in the ventral cross-section view of wild type and *shi^ts^* embryos during ventral furrow formation. Bottom panels (D’): En face view images showing myosin accumulation at the indicated time points. En face view images were Gaussian filtered with a radius of 0.5 pixel. Contrasts were adjusted to make the cytoplasmic background comparable. Scale bars, 10 μm. Embryos were placed at restrictive temperature (32°C) during cellularization. *shi^ts^* mutant embryos (N = 17 embryos) exhibit ventral furrow formation defects with different level of severity compared to wildtype controls (N = 5 embryos). *shi^ts^* (mild) embryos (N = 10 embryos) progress slower in ventral furrow formation while *shi^ts^* (severe) embryos (N = 7 embryos) fail to form ventral furrow as tissue relaxes after initial constriction. In both group of mutant embryos, apical myosin intensity eventually rises to a level comparable or higher than that in the wildtype controls. Interestingly, a previous study has shown that blocking endocytosis using *shi^ts^* in *snail* mutant embryos rescued the accumulation of Fog in the apical cytoplasm and partially restored apical Myosin II accumulation (Pouille et al., 2009). It is unclear whether the Myosin II phenotype observed in our study was associated with the role of dynamin in regulating Fog. **(E, E’)** Disruption of dynamin function does not prevent apical accumulation of Rab11 vesicles. (E) En face view (maximum projection over 2 μm from apical surface) of the ventral region of embryos at the onset of gastrulation (T = 0) or during gastrulation when ventral furrow shows similar morphology between different groups of embryos. (E’) Zoom-in view of the highlighted regions (red boxes) are shown at the bottom. *shi^ts^* (mild) embryos show similar apical Rab11 vesicle accumulation compared to the controls. In *shi^ts^* (severe) embryos, Rab11 vesicles are already present in the broad apical region of the embryo before apical constriction starts. Nevertheless, the amount of apical Rab11 vesicles substantially increases in the constricting cells during ventral furrow formation. Contrasts were adjusted to make the cytoplasmic background comparable. Images were Gaussian filtered with a radius of 0.7 pixel. Scale bar, 10 μm. Scale bar for zoom-in panel, 2 μm.

**Figure S3.**
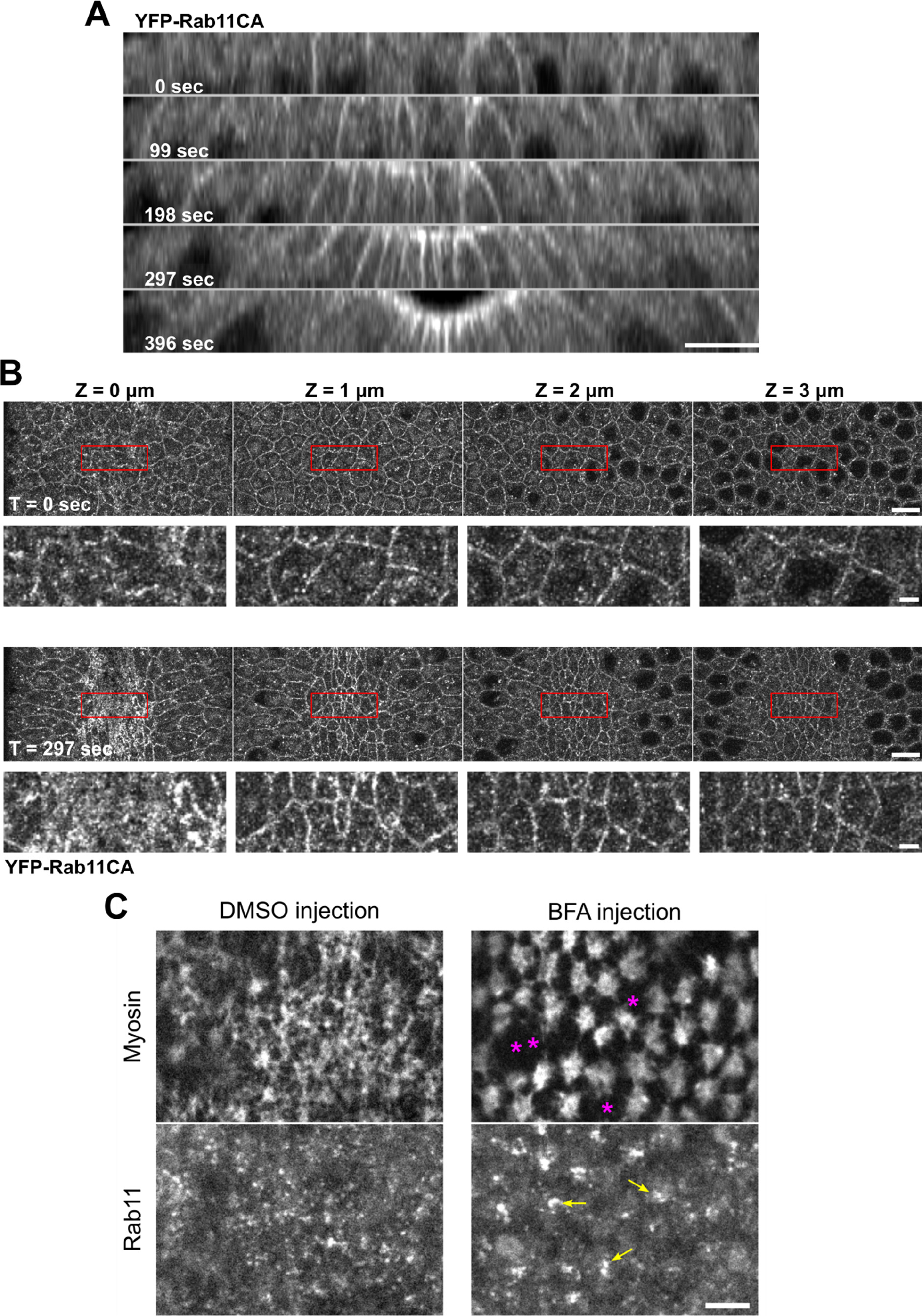
Rab11 vesicles are likely involved in the exocytic pathways. **(A, B)** Constitutively active Rab11 localizes to the plasma membrane in addition to vesicles and perinuclear compartments. (A) Montage showing the cross-section view of an embryo expressing YFP-Rab11CA during early ventral furrow formation. T = 0 sec indicates the onset of apical constriction. Apical and lateral membrane localization of Rab11CA is enhanced over time in the constricting cells. N = 4 embryos. Scale bar, 10 μm. (B) En face view of the same embryo at T = 0 sec and 297 sec from the apical surface (0 μm) to 4 μm below the surface. Scale bar, 10 μm. Bottom panels for each time point shows enlarged view of regions indicated by red boxes. Scale bar, 2 μm. **(C)** Acute inhibition of secretory pathway by Brefeldin A injection results in enlarged Rab11 structures (yellow arrows) compared to control DMSO injection. Breaks between neighboring myosin (magenta asterisks) was often observed in BFA injected embryos. N = 3 embryos. Scale bar, 5 μm.

**Figure S4.**
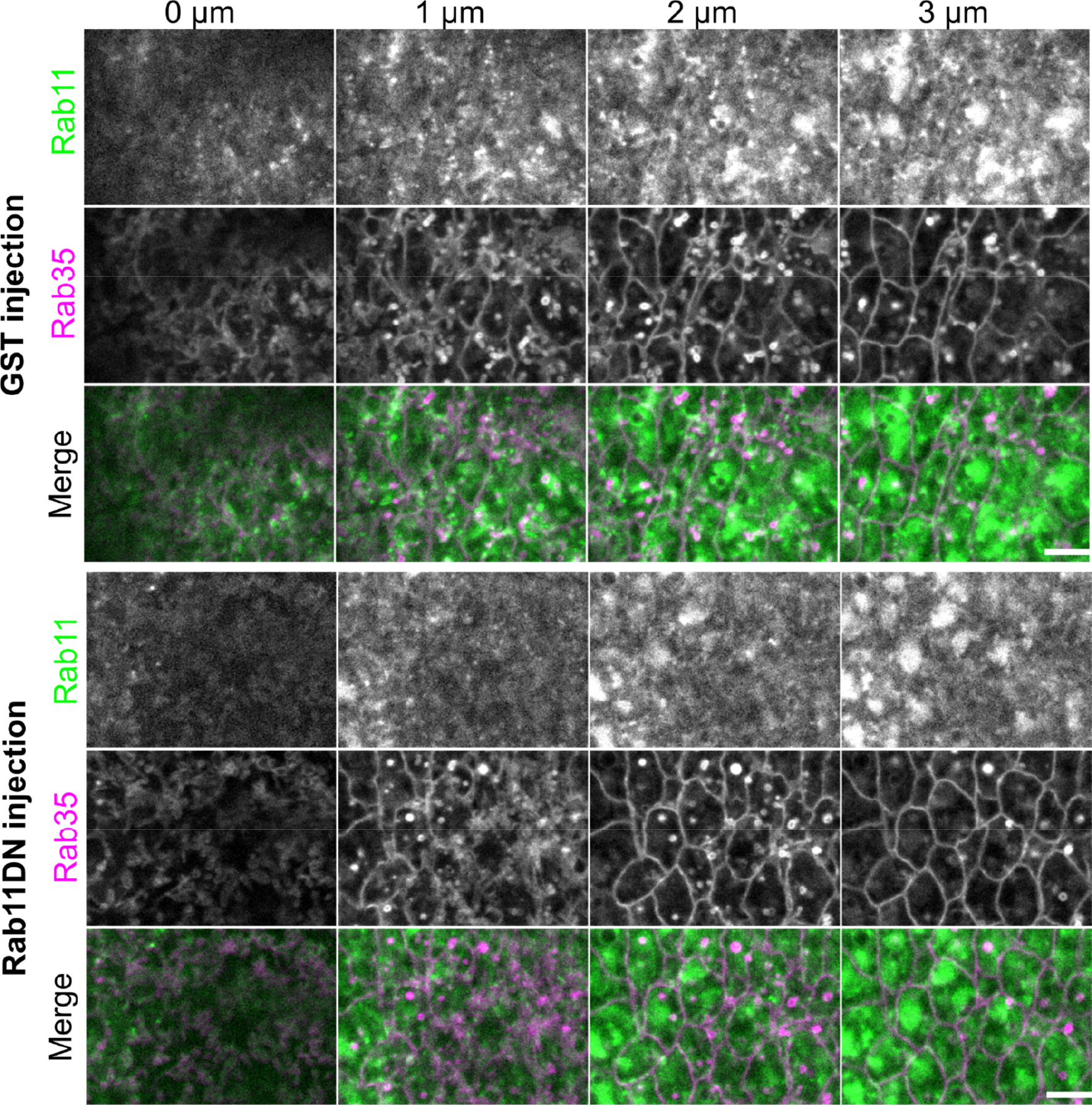
Rab11DN injection does not affect the Rab35 compartments during apical constriction. Eliminating apical Rab11 vesicle accumulation by Rab11DN injection does not prevent the formation of Rab35 positive tubules or obviously affect their morphology in the constricting cells during ventral furrow formation. Scale bar, 5 μm.

**Figure S5.**
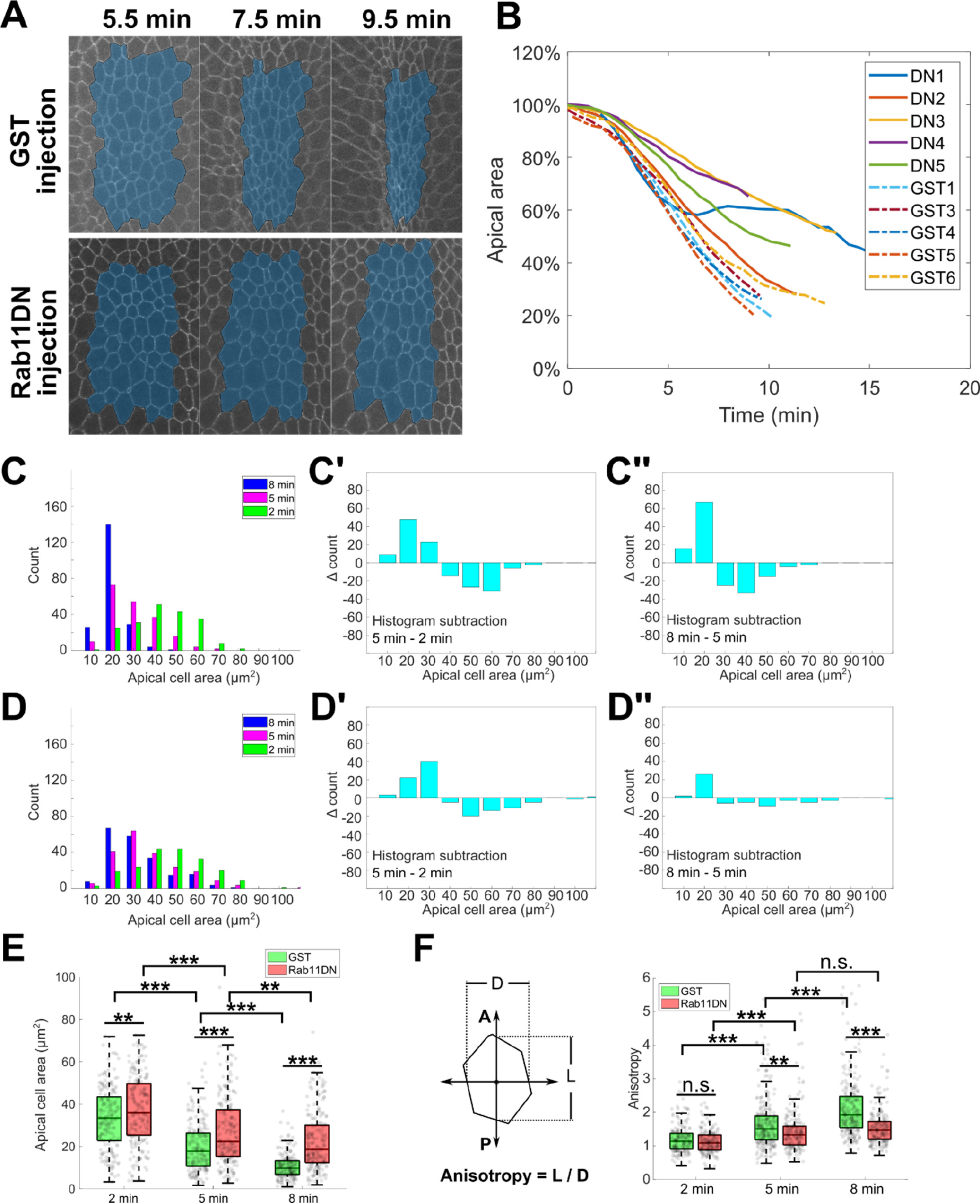
Acute inhibition of Rab11 results in reduced rate of apical constriction. **(A)** Representative images showing apical cell area change in GST injected embryo and Rab11DN injected embryo during apical constriction. A single z plane at 3-4 μm depth from the apical surface is shown. The total area of a cell group (masked in blue) located in the middle of the constriction domain was tracked over time and quantified in (B). **(B)** Apical area change over time. 5 embryos were injected with Rab11DN (DN1-DN5) and 6 embryos were injected with GST (GST1-GST6). The GST2 embryo was not used for quantification because it exhibited abnormal cell shape even before gastrulation. **(C-D)** Histograms of individual cell area distribution at 2min, 5min and 8min after the onset of gastrulation for GST (C, N = 4 embryos, GST1 embryo was not included due to lack of data at 2 min) and Rab11DN (D, N = 5 embryos) injected embryos. The change in area distribution is reflected by histogram subtraction between neighboring time points in the corresponding background (C’, C’’, D’, D’’). In the 2 – 5 minute period, apical area reduction in GST (C’) and Rab11DN (D’) injected embryos is in general comparable. However, in the 5 – 8 minute period, while cells in GST (C’’) injected embryos continue to reduce their apical area, cells in Rab11DN (D’’) injected embryos only show a very mild area reduction. **(E)** Box plot showing the average cell area at indicated time point for GST and Rab11DN injected embryos. Two tailed Student’s t-test was used for statistical comparison. **(F)** Quantification of anisotropy for individual constricting cells at 2 min, 5 min and 8 min after the onset of gastrulation. Same embryos from C-E were analyzed. Compared to GST injected control embryos, Rab11DN injected embryos show a substantially attenuated increase in anisotropy in the 5 – 8 minute period. Error bars stand for standard deviation. Two tailed Student’s t-test was used for statistical comparison.

## Supplemental Videos

**Video 1. Acute cytoD injection results in rapid tissue relaxation without immediately eliminating apical Myosin II.** 5 mg/mL cytoD was injected into embryo after onset of apical constriction. A maximum intensity projection of 3 μm of Myosin II signal is shown. Images were acquired every 9 s and the video is displayed at 10 frames per second.

**Video 2. Endocytosed FM4-64 vesicle colocalizes with Rab5 vesicle but not Rab11 vesicles.** 0.5 μg/μL FM4-64 was injected into the embryo at late cellularization. Images were acquired every 0.46 s during apical constriction and videos are displayed at 20 frames per second.

**Video 3. Acute disruption of MTs inhibits Rab11 vesicle transport.** A representative embryo immediately before (top) or one minute after (bottom) 25 mM colchicine injection is shown. Images were acquired every 0.22 s during apical constriction and videos are displayed at 20 frames per second.

**Video 4. Acute inactivation of Dynein inhibits Rab11 vesicle transport.** Top, Rab11 channel at indicated time after antibody injection; bottom, same corresponding movies overlaid with manually segmented trajectories, with red being apically directed transport and green being basally directed transport. 2 mg/mL antibody was injected during apical constriction. Scale bar, 2 μm. Images were acquired every 0.22 s during apical constriction and videos are displayed at 20 frames per second.

**Video 5. Rab11 vesicle dynamics near apical adherens junctions (marked by E-cadherin-GFP).** Rab11 vesicles are moving towards and become enriched at adherens junctions. Scale bar, 4 μm. Images were acquired every 0.21 s during apical constriction and videos are displayed at 20 frames per second.

**Video 6. Rab11 vesicles disappear after Rab11DN injection.** Maximum intensity projections of 3 μm Z stacks from apical surface. GST injection does not affect apical Rab11 vesicle accumulation. Upon Rab11DN injection, however, the apically accumulated Rab11 vesicles disappear over a 4-minute period, while the perinuclear Rab11 compartments appear to be intact. An exponential fit pixel-wise bleach correction method (ImageJ) was used to correct for photobleaching. Images were acquired every 5.8 s (left) or 7.2 s during apical constriction and videos are displayed at 10 frames per second.

**Video 7. Injection of Rab11DN causes frequent breaks in supracellular actomyosin network.** First part of the movie: Increased breaks in apical myosin network. Stars mark the site where myosin break events happen. Images were acquired every 5.8 s (left) or 5 s (right) during apical constriction and videos are displayed at 5 frames per second. Second part of the movie: Increased breaks in apical F-actin networks. Images were acquired every 6 s (top) or 7 s (bottom) during apical constriction and videos are displayed at 5 frames per second.

**Video 8. Acute dynein inhibition by dynein antibody injection affects apical Rab11 vesicle accumulation and causes frequent Myosin II breaks.** Dynein antibody was injected around onset of apical constriction or slightly after. Images were acquired every 4.1 s (top) or 4.7 s (bottom) during apical constriction and videos are displayed at 5 frames per second.

**Video 9. Brefeldin A injection results in abnormal Rab11 structures and frequent Myosin II breaks.** BFA was injected around late cellularization. Images were acquired every 30 s during apical constriction and videos are displayed at 2 frames per second.

**Video 10. Knockdown of Canoe causes frequent Myosin II breaks.** Images were acquired every 5 s and the video is displayed at 10 frames per second.

## References

1. Ahmed, W.W., and Saif, T.A. (2014). Active transport of vesicles in neurons is modulated by mechanical tension. Scientific Reports 4, 4481.

2. Ahmed, W.W., Li, T.C., Rubakhin, S.S., Chiba, A., Sweedler, J.V., and Saif, T.A. (2012). Mechanical Tension Modulates Local and Global Vesicle Dynamics in Neurons. Cel. Mol. Bioeng. 5, 155–164.

3. Amano, M., Ito, M., Kimura, K., Fukata, Y., Chihara, K., Nakano, T., Matsuura, Y., and Kaibuchi, K. (1996). Phosphorylation and Activation of Myosin by Rho-associated Kinase (Rho-kinase)*. Journal of Biological Chemistry 271, 20246–20249.

4. Benli, M., Döring, F., Robinson, D.G., Yang, X., and Gallwitz, D. (1996). Two GTPase isoforms, Ypt31p and Ypt32p, are essential for Golgi function in yeast. The EMBO Journal 15, 6460–6475.

5. Berg, S., Kutra, D., Kroeger, T., Straehle, C.N., Kausler, B.X., Haubold, C., Schiegg, M., Ales, J., Beier, T., Rudy, M., et al. (2019). ilastik: interactive machine learning for (bio)image analysis. Nat Methods 16, 1226–1232.

6. van der Bliek, A.M., and Meyerowrtz, E.M. (1991). Dynamin-like protein encoded by the Drosophila shibire gene associated with vesicular traffic. Nature 351, 411–414.

7. Boycott, H.E., Barbier, C.S.M., Eichel, C.A., Costa, K.D., Martins, R.P., Louault, F., Dilanian, G., Coulombe, A., Hatem, S.N., and Balse, E. (2013). Shear stress triggers insertion of voltage-gated potassium channels from intracellular compartments in atrial myocytes. Proceedings of the National Academy of Sciences 110, E3955–E3964.

8. Chanet, S., Miller, C.J., Vaishnav, E.D., Ermentrout, B., Davidson, L.A., and Martin, A.C. (2017). Actomyosin meshwork mechanosensing enables tissue shape to orient cell force. Nat Commun 8.

9. Chardin, P., and McCormick, F. (1999). Brefeldin A: The Advantage of Being Uncompetitive. Cell 97, 153–155.

10. Chen, W., Feng, Y., Chen, D., and Wandinger-Ness, A. (1998). Rab11 Is Required for Trans-Golgi Network–to–Plasma Membrane Transport and a Preferential Target for GDP Dissociation Inhibitor. Mol Biol Cell 9, 3241–3257.

11. Collinet, C., and Lecuit, T. (2021). Programmed and self-organized flow of information during morphogenesis. Nature Reviews Molecular Cell Biology 1–21.

12. Coravos, J.S., and Martin, A.C. (2016). Apical Sarcomere-like Actomyosin Contracts Nonmuscle Drosophila Epithelial Cells. Developmental Cell 39, 346–358.

13. Dawes-Hoang, R.E., Parmar, K.M., Christiansen, A.E., Phelps, C.B., Brand, A.H., and Wieschaus, E.F. (2005). folded gastrulation, cell shape change and the control of myosin localization. Development 132, 4165–4178.

14. Denk-Lobnig, M., Totz, J.F., Heer, N.C., Dunkel, J., and Martin, A.C. (2021). Combinatorial patterns of graded RhoA activation and uniform F-actin depletion promote tissue curvature. Development 148, dev199232.

15. Dillman, J.F., and Pfister, K.K. (1994). Differential phosphorylation in vivo of cytoplasmic dynein associated with anterogradely moving organelles. J Cell Biol 127, 1671–1681.

16. Dunst, S., Kazimiers, T., von Zadow, F., Jambor, H., Sagner, A., Brankatschk, B., Mahmoud, A., Spannl, S., Tomancak, P., Eaton, S., et al. (2015). Endogenously Tagged Rab Proteins: A Resource to Study Membrane Trafficking in Drosophila. Developmental Cell 33, 351–365.

17. Fletcher, D.A., and Mullins, R.D. (2010). Cell mechanics and the cytoskeleton. Nature 463, 485–492.

18. Fu, M., and Holzbaur, E.L.F. (2013). JIP1 regulates the directionality of APP axonal transport by coordinating kinesin and dynein motors. J Cell Biol 202, 495–508.

19. Fuse, N., Yu, F., and Hirose, S. (2013). Gprk2 adjusts Fog signaling to organize cell movements in Drosophila gastrulation. Development 140, 4246–4255.

20. Gauthier, N.C., Fardin, M.A., Roca-Cusachs, P., and Sheetz, M.P. (2011). Temporary increase in plasma membrane tension coordinates the activation of exocytosis and contraction during cell spreading. Proceedings of the National Academy of Sciences 108, 14467–14472.

21. Gelbart, M.A., He, B., Martin, A.C., Thiberge, S.Y., Wieschaus, E.F., and Kaschube, M. (2012). Volume conservation principle involved in cell lengthening and nucleus movement during tissue morphogenesis. Proceedings of the National Academy of Sciences 109, 19298–19303.

22. Gheisari, E., Aakhte, M., and Müller, H.-A.J. (2020). Gastrulation in Drosophila melanogaster: Genetic control, cellular basis and biomechanics. Mechanisms of Development 163, 103629.

23. Gilmour, D., Rembold, M., and Leptin, M. (2017). From morphogen to morphogenesis and back. Nature 541, 311–320.

24. Groth, A.C., Fish, M., Nusse, R., and Calos, M.P. (2004). Construction of Transgenic Drosophila by Using the Site-Specific Integrase From Phage φC31. Genetics 166, 1775–1782.

25. Grünfelder, C.G., Engstler, M., Weise, F., Schwarz, H., Stierhof, Y.-D., Morgan, G.W., Field, M.C., and Overath, P. (2003). Endocytosis of a Glycosylphosphatidylinositol-anchored Protein via Clathrin-coated Vesicles, Sorting by Default in Endosomes, and Exocytosis via RAB11-positive Carriers. MBoC 14, 2029–2040.

26. Guglielmi, G., Barry, J.D., Huber, W., and De Renzis, S. (2015). An Optogenetic Method to Modulate Cell Contractility during Tissue Morphogenesis. Developmental Cell 35, 646–660.

27. Guo, H., Swan, M., Huang, S., and He, B. (2021). Mechanical bistability enabled by ectodermal compression facilitates Drosophila mesoderm invagination.

28. He, B., Doubrovinski, K., Polyakov, O., and Wieschaus, E. (2014). Apical constriction drives tissue-scale hydrodynamic flow to mediate cell elongation. Nature 508, 392.

29. Heer, N.C., Miller, P.W., Chanet, S., Stoop, N., Dunkel, J., and Martin, A.C. (2017). Actomyosin-based tissue folding requires a multicellular myosin gradient. Development 144, 1876–1886.

30. Horgan, C.P., Hanscom, S.R., Jolly, R.S., Futter, C.E., and McCaffrey, M.W. (2010). Rab11- FIP3 links the Rab11 GTPase and cytoplasmic dynein to mediate transport to the endosomal- recycling compartment. Journal of Cell Science 123, 181–191.

31. Jedd, G., Mulholland, J., and Segev, N. (1997). Two New Ypt GTPases Are Required for Exit From the Yeast trans-Golgi Compartment. J Cell Biol 137, 563–580.

32. Jewett, C.E., Vanderleest, T.E., Miao, H., Xie, Y., Madhu, R., Loerke, D., and Blankenship, J.T. (2017). Planar polarized Rab35 functions as an oscillatory ratchet during cell intercalation in the Drosophila epithelium. Nature Communications 8, 476.

33. Jha, A., van Zanten, T.S., Philippe, J.-M., Mayor, S., and Lecuit, T. (2018). Quantitative Control of GPCR Organization and Signaling by Endocytosis in Epithelial Morphogenesis. Current Biology 28, 1570–1584.e6.

34. Jodoin, J.N., Coravos, J.S., Chanet, S., Vasquez, C.G., Tworoger, M., Kingston, E.R., Perkins, L.A., Perrimon, N., and Martin, A.C. (2015). Stable Force Balance between Epithelial Cells Arises from F-Actin Turnover. Developmental Cell 35, 685–697.

35. Karpova, N., Bobinnec, Y., Fouix, S., Huitorel, P., and Debec, A. (2006). Jupiter, a new Drosophila protein associated with microtubules. Cell Motility 63, 301–312.

36. Kennedy, M.J., Hughes, R.M., Peteya, L.A., Schwartz, J.W., Ehlers, M.D., and Tucker, C.L. (2010). Rapid blue-light–mediated induction of protein interactions in living cells. Nature Methods 7, 973–975.

37. Kerridge, S., Munjal, A., Philippe, J.-M., Jha, A., de las Bayonas, A.G., Saurin, A.J., and Lecuit, T. (2016). Modular activation of Rho1 by GPCR signalling imparts polarized myosin II activation during morphogenesis. Nature Cell Biology 18, 261–270.

38. Khandelwal, P., Prakasam, H.S., Clayton, D.R., Ruiz, W.G., Gallo, L.I., van Roekel, D., Lukianov, S., Peränen, J., Goldenring, J.R., and Apodaca, G. (2013). A Rab11a-Rab8a-Myo5B network promotes stretch-regulated exocytosis in bladder umbrella cells. MBoC 24, 1007–1019.

40. Kimura, K., Ito, M., Amano, M., Chihara, K., Fukata, Y., Nakafuku, M., Yamamori, B., Feng, J., Nakano, T., Okawa, K., et al. (1996). Regulation of Myosin Phosphatase by Rho and Rho- Associated Kinase (Rho-Kinase). Science 273, 245–248.

41. Kirby, T.J., and Lammerding, J. (2016). Stretch to express. Nature Materials 15, 1227–1229.

42. Ko, C.S., Tserunyan, V., and Martin, A.C. (2019). Microtubules promote intercellular contractile force transmission during tissue folding. J Cell Biol 218, 2726–2742.

43. Langevin, J., Morgan, M.J., Rossé, C., Racine, V., Sibarita, J.-B., Aresta, S., Murthy, M., Schwarz, T., Camonis, J., and Bellaïche, Y. (2005). Drosophila Exocyst Components Sec5, Sec6, and Sec15 Regulate DE-Cadherin Trafficking from Recycling Endosomes to the Plasma Membrane. Developmental Cell 9, 365–376.

44. Lapierre, L.A., Kumar, R., Hales, C.M., Navarre, J., Bhartur, S.G., Burnette, J.O., Provance, D.W., Mercer, J.A., Bähler, M., and Goldenring, J.R. (2001). Myosin Vb Is Associated with Plasma Membrane Recycling Systems. MBoC 12, 1843–1857.

45. Le, T.P., and Chung, S. (2021). Regulation of apical constriction via microtubule- and Rab11- dependent apical transport during tissue invagination. BioRxiv 827378.

46. Le Droguen, P.-M., Claret, S., Guichet, A., and Brodu, V. (2015). Microtubule-dependent apical restriction of recycling endosomes sustains adherens junctions during morphogenesis of the Drosophila tracheal system. Development 142, 363–374.

47. Lee, J.-Y., and Harland, R.M. (2010). Endocytosis Is Required for Efficient Apical Constriction during Xenopus Gastrulation. Current Biology 20, 253–258.

48. Leptin, M., and Grunewald, B. (1990). Cell shape changes during gastrulation in Drosophila. Development 110, 73–84.

49. Ligoxygakis, P., Roth, S., and Reichhart, J.-M. (2003). A Serpin Regulates Dorsal-Ventral Axis Formation in the Drosophila Embryo. Current Biology 13, 2097–2102.

50. Lim, B., Levine, M., and Yamazaki, Y. (2017). Transcriptional Pre-patterning of Drosophila Gastrulation. Current Biology 27, 286–290.

51. Lipatova, Z., Tokarev, A.A., Jin, Y., Mulholland, J., Weisman, L.S., and Segev, N. (2008). Direct Interaction between a Myosin V Motor and the Rab GTPases Ypt31/32 Is Required for Polarized Secretion. MBoC 19, 4177–4187.

52. Lock, J.G., and Stow, J.L. (2005). Rab11 in Recycling Endosomes Regulates the Sorting and Basolateral Transport of E-Cadherin. MBoC 16, 1744–1755.

53. Manning, A.J., Peters, K.A., Peifer, M., and Rogers, S.L. (2013). Regulation of Epithelial Morphogenesis by the G Protein–Coupled Receptor Mist and Its Ligand Fog. Sci. Signal. 6, ra98–ra98.

54. Martin, A.C. (2020). The Physical Mechanisms of Drosophila Gastrulation: Mesoderm and Endoderm Invagination. Genetics 214, 543–560.

55. Martin, A.C., and Goldstein, B. (2014). Apical constriction: themes and variations on a cellular mechanism driving morphogenesis. Development 141, 1987–1998.

56. Martin, A.C., Gelbart, M., Fernandez-Gonzalez, R., Kaschube, M., and Wieschaus, E.F. (2010). Integration of contractile forces during tissue invagination. The Journal of Cell Biology 188, 735–749.

57. Martin AC, Kaschube M, Wieschaus EF. Pulsed contractions of an actin-myosin network drive apical constriction. Nature 2009;457:495–9.

58. Mason, F.M., Tworoger, M., and Martin, A.C. (2013). Apical domain polarization localizes actin–myosin activity to drive ratchet-like apical constriction. Nat Cell Biol 15, 926–936.

59. Mateus, A.M., Gorfinkiel, N., Schamberg, S., and Martinez Arias, A. (2011). Endocytic and Recycling Endosomes Modulate Cell Shape Changes and Tissue Behaviour during Morphogenesis in Drosophila. PLoS One 6.

60. Mazumdar, A., and Mazumdar, M. (2002). How one becomes many: Blastoderm cellularization in Drosophila melanogaster. BioEssays 24, 1012–1022.

61. Miao, H., Vanderleest, T.E., Jewett, C.E., Loerke, D., and Blankenship, J.T. (2019). Cell ratcheting through the Sbf RabGEF directs force balancing and stepped apical constriction. J Cell Biol 218, 3845–3860.

62. Morize, P., Christiansen, A.E., Costa, M., Parks, S., and Wieschaus, E. (1998). Hyperactivation of the folded gastrulation pathway induces specific cell shape changes. Development 125, 589– 597.

63. Moughamian, A.J., Osborn, G.E., Lazarus, J.E., Maday, S., and Holzbaur, E.L.F. (2013). Ordered Recruitment of Dynactin to the Microtubule Plus-End is Required for Efficient Initiation of Retrograde Axonal Transport. Journal of Neuroscience 33, 13190–13203.

64. Narumiya, S., Ishizaki, T., and Ufhata, M. (2000). Use and properties of ROCK-specific inhibitor Y-27632. In Methods in Enzymology, W.E. Balch, C.J. Der, and A. Hall, eds. (Academic Press), pp. 273–284.

65. Nikolaidou, K.K., and Barrett, K. (2004). A Rho GTPase Signaling Pathway Is Used Reiteratively in Epithelial Folding and Potentially Selects the Outcome of Rho Activation. Current Biology 14, 1822–1826.

66. Ollion, J., Cochennec, J., Loll, F., Escudé, C., and Boudier, T. (2013). TANGO: a generic tool for high-throughput 3D image analysis for studying nuclear organization. Bioinformatics 29, 1840–1841.

67. Ossipova, O., Kim, K., Lake, B.B., Itoh, K., Ioannou, A., and Sokol, S.Y. (2014). Role of Rab11 in planar cell polarity and apical constriction during vertebrate neural tube closure. Nature Communications 5, 3734.

68. Ossipova, O., Chuykin, I., Chu, C.-W., and Sokol, S.Y. (2015). Vangl2 cooperates with Rab11 and Myosin V to regulate apical constriction during vertebrate gastrulation. Development 142, 99–107.

69. Otani, T., Oshima, K., Onishi, S., Takeda, M., Shinmyozu, K., Yonemura, S., and Hayashi, S. (2011). IKKε regulates cell elongation through recycling endosome shuttling. Dev Cell 20, 219–232.

70. Pelissier, A., Chauvin, J.-P., and Lecuit, T. (2003). Trafficking through Rab11 Endosomes Is Required for Cellularization during Drosophila Embryogenesis. Current Biology 13, 1848– 1857.

71. Pouille, P.-A., Ahmadi, P., Brunet, A.-C., and Farge, E. (2009). Mechanical Signals Trigger Myosin II Redistribution and Mesoderm Invagination in Drosophila Embryos. Sci. Signal. 2, ra16–ra16.

72. Razzell, W., Bustillo, M.E., and Zallen, J.A. (2018). The force-sensitive protein Ajuba regulates cell adhesion during epithelial morphogenesis. J Cell Biol 217, 3715–3730.

73. Riggs, B., Rothwell, W., Mische, S., Hickson, G.R.X., Matheson, J., Hays, T.S., Gould, G.W., and Sullivan, W. (2003). Actin cytoskeleton remodeling during early Drosophila furrow formation requires recycling endosomal components Nuclear-fallout and Rab11. Journal of Cell Biology 163, 143–154.

74. Sawyer, J.K., Harris, N.J., Slep, K.C., Gaul, U., and Peifer, M. (2009). The Drosophila afadin homologue Canoe regulates linkage of the actin cytoskeleton to adherens junctions during apical constriction. Journal of Cell Biology 186, 57–73.

75. Schonteich, E., Wilson, G.M., Burden, J., Hopkins, C.R., Anderson, K., Goldenring, J.R., and Prekeris, R. (2008). The Rip11/Rab11-FIP5 and kinesin II complex regulates endocytic protein recycling. Journal of Cell Science 121, 3824–3833.

76. Schuh, M. (2011). An actin-dependent mechanism for long-range vesicle transport. Nature Cell Biology 13, 1431–1436.

77. Shillcock, J.C., and Lipowsky, R. (2005). Tension-induced fusion of bilayer membranes and vesicles. Nature Materials 4, 225–228.

78. Siechen, S., Yang, S., Chiba, A., and Saif, T. (2009). Mechanical tension contributes to clustering of neurotransmitter vesicles at presynaptic terminals. Proceedings of the National Academy of Sciences 106, 12611–12616.

79. Staykova, M., Holmes, D.P., Read, C., and Stone, H.A. (2011). Mechanics of surface area regulation in cells examined with confined lipid membranes. PNAS 108, 9084–9088.

80. Sun, S., and Irvine, K.D. (2016). Cellular Organization and Cytoskeletal Regulation of the Hippo Signaling Network. Trends in Cell Biology 26, 694–704.

81. Sweeton, D., Parks, S., Costa, M., and Wieschaus, E. (1991). Gastrulation in Drosophila: the formation of the ventral furrow and posterior midgut invaginations. Development 112, 775– 789.

82. Takahashi, S., Kubo, K., Waguri, S., Yabashi, A., Shin, H.-W., Katoh, Y., and Nakayama, K. (2012). Rab11 regulates exocytosis of recycling vesicles at the plasma membrane. J Cell Sci 125, 4049–4057.

83. Tiwari, A.K., and Roy, J.K. (2008). Rab11 is essential for fertility in Drosophila. Cell Biology International 32, 1158–1168.

84. Uhler, C., and Shivashankar, G.V. (2017). Regulation of genome organization and gene expression by nuclear mechanotransduction. Nature Reviews Molecular Cell Biology 18, 717– 727.

85. Ullrich, O., Reinsch, S., Urbé, S., Zerial, M., and Parton, R.G. (1996). Rab11 regulates recycling through the pericentriolar recycling endosome. J Cell Biol 135, 913–924.

86. Vasquez, C.G., Tworoger, M., and Martin, A.C. (2014). Dynamic myosin phosphorylation regulates contractile pulses and tissue integrity during epithelial morphogenesis. J Cell Biol 206, 435–450.

87. Vaughan, P.S., Miura, P., Henderson, M., Byrne, B., and Vaughan, K.T. (2002). A role for regulated binding of p150 *^Glued^* to microtubule plus ends in organelle transport. The Journal of Cell Biology 158, 305–319.

88. Wang, Y., Huynh, W., Skokan, T.D., Lu, W., Weiss, A., and Vale, R.D. (2019). CRACR2a is a calcium-activated dynein adaptor protein that regulates endocytic traffic. The Journal of Cell Biology jcb.201806097.

89. Wang, Y.-C., Khan, Z., Kaschube, M., and Wieschaus, E.F. (2012). Differential positioning of adherens junctions is associated with initiation of epithelial folding. Nature 484, 390–393.

90. Wang, Z., Edwards, J.G., Riley, N., Provance, D.W., Karcher, R., Li, X., Davison, I.G., Ikebe, M., Mercer, J.A., Kauer, J.A., et al. (2008). Myosin Vb Mobilizes Recycling Endosomes and AMPA Receptors for Postsynaptic Plasticity. Cell 135, 535–548.

91. Welz, T., Wellbourne-Wood, J., and Kerkhoff, E. (2014). Orchestration of cell surface proteins by Rab11. Trends in Cell Biology 24, 407–415.

92. Weng, M., and Wieschaus, E. (2016). Myosin-dependent remodeling of adherens junctions protects junctions from Snail-dependent disassembly. Journal of Cell Biology 212, 219–229.

93. Winter, C.G., Wang, B., Ballew, A., Royou, A., Karess, R., Axelrod, J.D., and Luo, L. (2001). Drosophila Rho-Associated Kinase (Drok) Links Frizzled-Mediated Planar Cell Polarity Signaling to the Actin Cytoskeleton. Cell 105, 81–91.

94. Woichansky, I., Beretta, C.A., Berns, N., and Riechmann, V. (2016). Three mechanisms control E-cadherin localization to the zonula adherens. Nature Communications 7.

95. Wu, X.-S., Elias, S., Liu, H., Heureaux, J., Wen, P.J., Liu, A.P., Kozlov, M.M., and Wu, L.-G. (2017). Membrane Tension Inhibits Rapid and Slow Endocytosis in Secretory Cells. Biophysical Journal 113, 2406–2414.

96. Yashiro, H., Loza, A.J., Skeath, J.B., and Longmore, G.D. (2014). Rho1 regulates adherens junction remodeling by promoting recycling endosome formation through activation of myosin II. MBoC 25, 2956–2969.

97. Yi, J.Y., Ori-McKenney, K.M., McKenney, R.J., Vershinin, M., Gross, S.P., and Vallee, R.B. (2011). High-resolution imaging reveals indirect coordination of opposite motors and a role for LIS1 in high-load axonal transport. Journal of Cell Biology 195, 193–201.

